# Principles of metabolome conservation in animals

**DOI:** 10.1101/2022.08.15.503737

**Authors:** Orsolya Liska, Gábor Boross, Charles Rocabert, Balázs Szappanos, Roland Tengölics, Balázs Papp

**Affiliations:** HCEMM-BRC Metabolic Systems Biology Lab; Szeged, Hungary; Synthetic and Systems Biology Unit, Institute of Biochemistry, Biological Research Centre, Eötvös Loránd Research Network (ELKH); Szeged, Hungary; Doctoral School of Biology, University of Szeged; Szeged, Hungary; Department of Biology, Stanford University, Stanford, California, USA; Inria, 78150 Rocquencourt, France; Organismal and Evolutionary Biology Research Programme, University of Helsinki, Helsinki, Finland; Department of Biotechnology, University of Szeged, Szeged, Hungary; Biological Research Centre, Metabolomics Lab, Eötvös Loránd Research Network (ELKH), Szeged, Hungary

**Author notes:** Corresponding authors: Balázs Papp and Gábor Boross. These authors have contributed equally to this work (Orsolya Liska, Gábor Boross, Charles Rocabert).

## Abstract

Metabolite concentrations shape cellular physiology and disease susceptibility, yet the general principles governing metabolome evolution are largely unknown. Here we introduce a measure of conservation of individual metabolite concentrations among related species. By analysing multispecies metabolome datasets in mammals and fruit flies, we show that conservation varies extensively across metabolites. Three major functional properties, metabolite abundance, essentiality and association with human diseases predict conservation, highlighting a striking parallel between the evolutionary forces driving metabolome and protein sequence conservation. Metabolic network simulations recapitulated these general patterns, and revealed that abundant metabolites are highly conserved due to their strong coupling to key metabolic fluxes in the network. This study uncovers simple rules governing metabolic evolution in animals and implies that most metabolome differences between species are permitted, rather than favored by selection. More broadly, our work paves the way towards using evolutionary information to discover biomarkers, as well as to detect pathogenic metabolome alterations in individual patients.

## Introduction

Metabolites are intermediates of biochemical pathways as well as regulators of enzymes and non-enzymatic proteins^1^. Accordingly, intracellular metabolite concentrations are key quantities that affect the rates of metabolic reactions (fluxes) and regulate various layers of cellular organization^2,3^. Consequently, metabolite dysregulation underlies various human diseases, from metabolic disorders to cancer^4^. Given the tight associations between metabolite concentrations and cellular physiology, it is often supposed that evolutionary changes in the metabolome contribute to phenotypic differences between species^5,6^. Indeed, shifts in specific metabolite concentrations have been associated with phenotypic evolution in both mammals^5,6^ and plants^7,8^. However, the general principles driving metabolome evolution remain largely unexplored. Metabolite concentrations show broad similarities between cells of distantly related organisms^9^ and obey simple optimality principles^10,11^, indicating widespread natural selection to preserve them. In fact, metabolome-altering mutations continuously occur during evolution, with harmful ones likely being eliminated by natural selection, affecting patterns of metabolome variation among species. Thus, elucidating the evolutionary forces shaping the metabolome has relevance for a better understanding of the organization of cellular metabolism, and for human health as well.

Here we propose that the functional role and biochemical properties of metabolites influence the amount of permissible concentration changes during evolution. Consequently, metabolites that are more strongly constrained by the requirement for proper cellular function, *i*.*e*. subject to stronger functional constraints, should be more evolutionarily conserved in their concentrations. This is analogous to the well-established phenomenon that proteins evolving under stronger functional constraints are more conserved in their sequences^12,13^. Furthermore, just as sequence conservation informs on deleterious genetic variants^14,15^, conservation of metabolite concentrations should inform on the health impact of metabolite changes. By combining phylogenetic analysis of metabolomics data with systems biology modelling, we found support for this hypothesis and revealed simple rules that dictate the evolution of metabolite concentrations in animals. Overall, our work offers a universal conceptual framework of metabolome conservation that informs on the disease association of metabolites.

## Results

### Extensive variation in evolutionary conservation of metabolite concentrations

To systematically investigate the evolutionary conservation of metabolite concentrations, we first focused on mammals due to their relevance for human health and the availability of comprehensive multispecies metabolomics data. We analyzed a published dataset that contains the relative concentrations of 139 non-lipid metabolites in four major organs (brain, kidney, heart and liver) across 26 mammalian species, spanning an evolutionary period of ∼200 million years^6^ (Fig. 1A). These metabolites cover a wide range of central metabolic pathways, including amino acid, carbohydrate, energy and cofactor metabolism.

**Figure 1:**
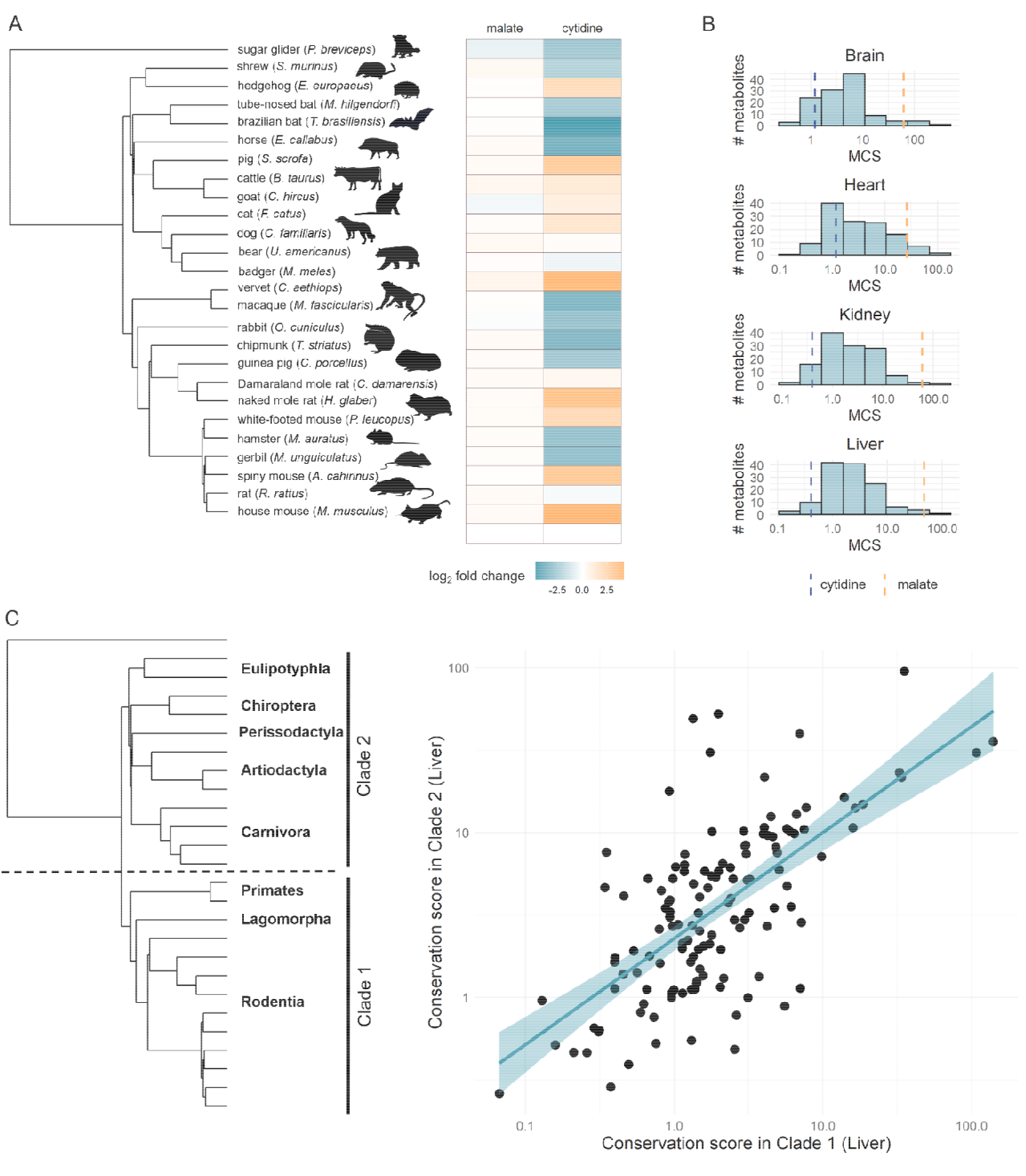
Metabolites differ widely in their conservation score during mammalian evolution. **A**: Phylogeny of 26 mammals and heatmap illustrating how metabolite levels in liver vary among species (log_2_ fold-change compared to mouse), exemplified by two metabolites: malate shows highly similar concentrations in all species, while cytidine is highly variable. **B**: Distributions of metabolite conservation scores in each organ, as inferred from all 26 species (logarithmic scale). The vertical dashed lines show the MCSs of the two example metabolites from panel A (cytidine – blue, malate – yellow). **C**: Conservation scores (liver) calculated for two independent clades of the tree show a strong correlation (Pearson’s r = 0.67, p = 1.1e-18, N = 132). Line depicts the fitted linear regression. Similar results are obtained for other organs (Suppl. Fig. S6). The tree on the left depicts the two independent clades of mammals for which conservation scores were inferred, with the names of the constituents’ taxonomical orders.

The magnitude of concentration differences among the 26 species varies widely across metabolites in all studied organs (Suppl. Fig. S1). For example, in liver, cytidine level varies up to 529-fold among species, while malate shows highly similar concentrations in all species, with less than 2.3-fold differences (Fig. 1A). This pattern suggests extensive variation in the evolutionary conservation of metabolite concentrations among different metabolites. To more rigorously estimate the degree of evolutionary conservation, while also accounting for species phylogeny, we introduce a score that captures the extent of conservation of metabolite concentrations along the phylogeny for each metabolite and for each organ (see Methods). This *metabolite conservation score* (MCS) is based on the widely used Brownian motion model of trait evolution^16^, and is defined as the inverse of the rate of evolutionary changes in metabolite concentration (see Methods).

MCS displays extensive variation across metabolites in all four organs, spanning 560-to 970-fold ranges (Fig. 1B). Importantly, variation in MCS is not caused by technical artefacts, as within-species variation and measurement noise explain less than ∼7% of total variance in MCS (Suppl. Fig. S2, Methods) and using an evolutionary model that explicitly incorporates such variation results in highly similar MCS values (Suppl. Fig. S3, Methods). Phylogenetic comparisons of metabolomes might be confounded by dietary and environmental differences between species. To address this issue, we additionally analysed a metabolome dataset of fibroblasts isolated from 16 mammals and cultured under identical *in vitro* conditions^17^. Despite large differences between cell types and study conditions, the MCS values calculated from the *in vitro* fibroblast data show highly significant correlations with those based on the *in vivo* assessment of the four organs (Suppl. Fig. S4). Furthermore, MCS extensively varies among different metabolites (up to 290-fold) in the fibroblast dataset as well. Thus, large differences in the conservation of individual metabolite concentrations hold in an independent dataset, and are not confounded by measurement noise or lifestyle differences.

Conservation of a metabolite might vary across the phylogeny due to lineage-specific shifts in selective pressure (*i*.*e*. adaptive evolution). However, such heterogeneities are unlikely to confound our inferences. First, in each organ, only a small fraction (7-29%) of metabolites show lineage-specific concentration shifts^6^ and MCS is robust to such effects (Suppl. Fig. S5, see Methods). Second, the conservation scores calculated on two independent clades of the tree — rodents, rabbit and primates versus all other species – strongly correlate for all organs (see Fig. 1C for liver and Suppl. Fig. S6 for other organs). Thus, metabolites that are conserved in a particular clade also tend to be conserved in the rest of mammals.

Together, these results indicate that the evolution of individual metabolite concentrations can be characterized by a simple conservation score which is largely invariant across mammalian clades, but varies extensively across metabolites.

### Abundance and essentiality are important determinants of evolutionary conservation

By analogy to the neutral theory of sequence evolution^18^, we hypothesize that conserved metabolites are subject to stronger functional constraints against concentration changes. If so, MCS should largely be determined by the functional role and biochemical properties of metabolites. To test this, we compiled 17 features capturing the biological and chemical properties of metabolites, including pathway membership, the number of network connections (network degree), absolute metabolite concentration (abundance), chemical class, toxicity, and various physicochemical properties (Fig. 2A and Suppl. Table S1, see Methods). We also included ‘essentiality’, a feature describing whether a metabolite participates in a reaction catalyzed by an essential enzyme, as determined by gene deletions in mice^19^.

**Figure 2:**
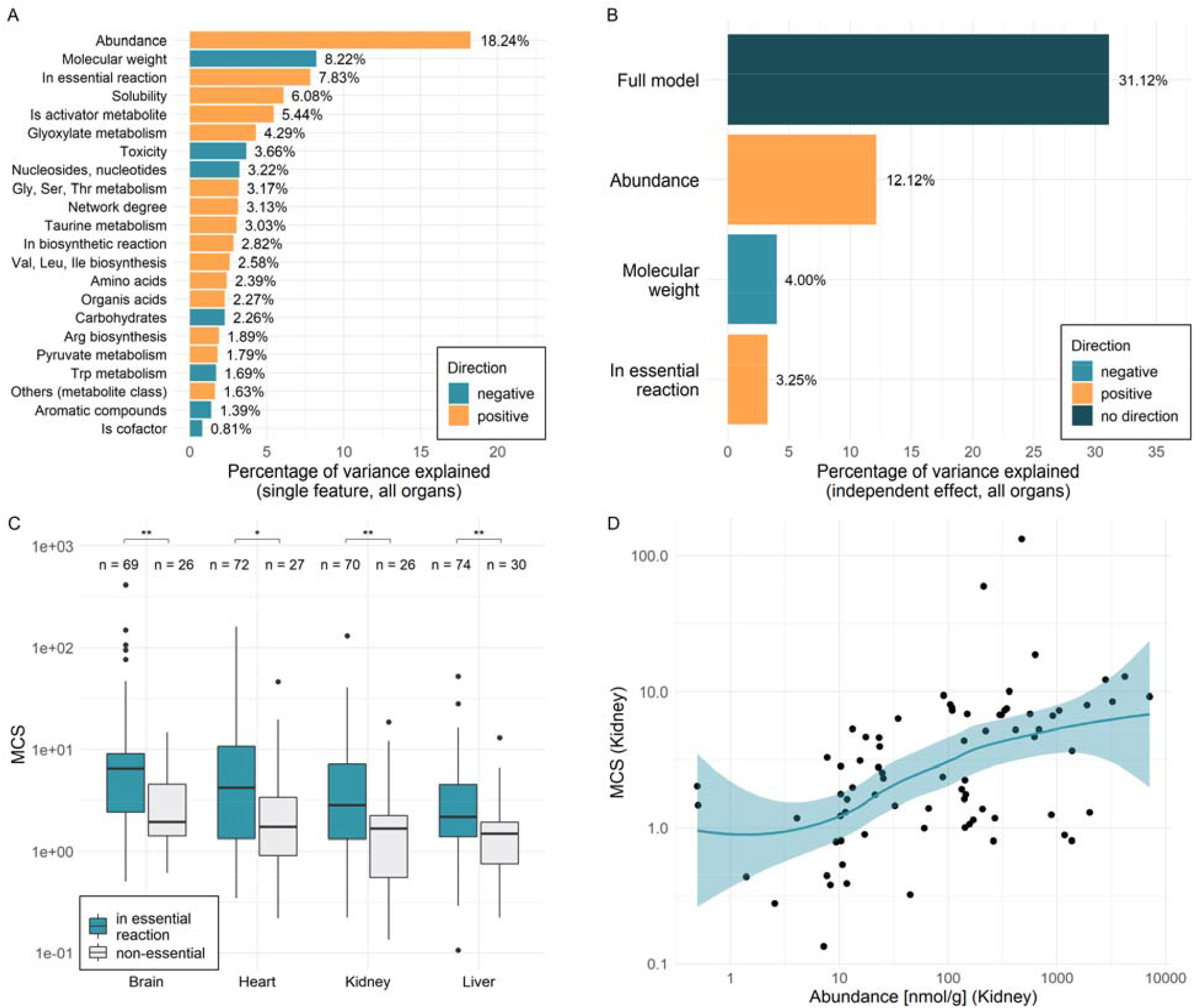
Determinants of metabolite conservation score. **A**: Barplot showing the percentage of variance in conservation score explained by each metabolite feature individually (i.e. univariate models). Only features with a significant effect are shown (FDR<0.01, Methods). **B**: Barplot showing the percentage of variance in conservation score explained by multivariate regression modelling. Bars indicate the percentage of variance explained by the full model, as well as the independent contribution of each feature found to be significant in the model (p<0.05). **C**: Metabolites participating in reactions encoded by essential genes show higher conservation scores than the rest of metabolites (two-sided Wilcoxon tests, brain: p = 0.0013; heart: p = 0.019; kidney p = 0.0049; liver p = 0.0022). Boxplots show the median, first and third quartiles, with the whiskers showing the values within a 1.5 interquartile range distance from the 1^st^ and 3^rd^ quartiles. **D:** Metabolites with high absolute concentration (abundance) show higher conservation scores (kidney: Spearman’s correlation rho = -0.53, p = 1.1e-06, N=76; for other organs see Suppl. Fig S7). Line indicates LOESS regression, with their 95% confidence interval indicated in blue.

We found several metabolite features that are statistically significantly associated with MCS when analysed individually (Fig. 2A, Suppl. Table S1, Methods). As metabolite features might correlate with each other, next we carried out multivariate regression to identify independent predictors of MCS by jointly analysing the four organs (Methods). We found that a simple model with three dominant predictors explains ∼31% of variation in MCS across metabolites (Fig. 2B, Suppl. Table S3). This is a remarkably high figure considering the inherent stochasticity of evolution. Indeed, we estimate an upper limit of 49% for the predictability of MCS based on its reproducibility between two independent clades (Fig 1C and Suppl. Table S2).

Three general metabolite properties have significant independent effects on MCS in all organs: abundance, molecular weight and involvement in essential reactions (Fig. 2B, Table S4). Specifically, metabolites that are (i) highly abundant in the cell, (ii) have a small molecular size or (iii) participate in reactions encoded by essential genes tend to be more conserved (Fig. 2C-D). The latter finding is consistent with the intuitive expectation that perturbations in metabolites associated with essential enzymes are especially harmful, and are therefore subject to stronger stabilizing selection.

Abundance is the strongest determinant of conservation in all organs (Fig. 2B, Suppl. Table S4). Metabolites vastly differ in their absolute concentrations, spanning over six orders of magnitude^9^. We revealed that higher abundance of a metabolite is associated with a higher conservation score, with a continuous trend across the entire range of abundance (Fig. 2D and Suppl. Fig. S7). We emphasize that abundance data comes from an independent study that quantified absolute metabolite concentrations for several organs in mice^20^. Low-abundance metabolites are typically measured with larger noise, potentially underestimating their conservation scores. However, such a bias is unlikely to confound our results because the positive relationship between abundance and MCS holds when explicitly accounting for measurement variability in the conservation score calculations (Supp. Table S5, Methods). Furthermore, the relationship also holds when inferring MCS from the *in vitro* fibroblast dataset, suggesting that it is not an artefact of comparing species with different diets (Suppl. Fig. S8).

Conservation is largely independent of the chemical class, physico-chemical properties, toxicity, pathway membership, network position and interaction degree of metabolites in the multivariate model (Suppl. Table S4). For instance, while metabolites participating in many reactions or serving as allosteric activators of enzymes tend to be conserved, these relationships disappear when accounting for other metabolite properties (Suppl. Table S4). Thus, contrary to intuitive expectations, metabolites interacting with multiple enzymes are not more strongly constrained. Furthermore, the toxic effects of highly increased metabolite levels, as assessed by toxicity in mice^21^, do not appear to constrain metabolome evolution.

Together, these findings demonstrate that evolutionary conservation of metabolite concentrations is well predictable based on a handful of metabolite properties, with abundance being the primary determinant.

### Organ-specific metabolite conservation reflects differential functional constraints

We next sought to investigate the differences in individual metabolite conservation scores between different organs. We hypothesize that such shifts in conservation arise from organ-specific biological functions.

In general, MCS correlates well among the four organs, indicating similar functional constraints (Fig. 3A). This is consistent with our finding that a handful of metabolite properties universally determine conservation in all four organs (Fig. 2B). Nevertheless, some metabolites are much more conserved in one organ than in the others (Suppl. Table S6). Literature data suggest that such differences partly reflect organ-specific functions (Fig. 3B). For example, both gamma-aminobutyrate (GABA) and glutamate show the strongest conservation in the brain, where they serve as the principal inhibitory and excitatory neurotransmitters, respectively^22^. Similarly, the osmolytes betaine and myo-inositol are especially conserved in the kidney and such molecules have key roles in protecting renal medullary cells from high NaCl and urea levels^23^.

**Figure 3:**
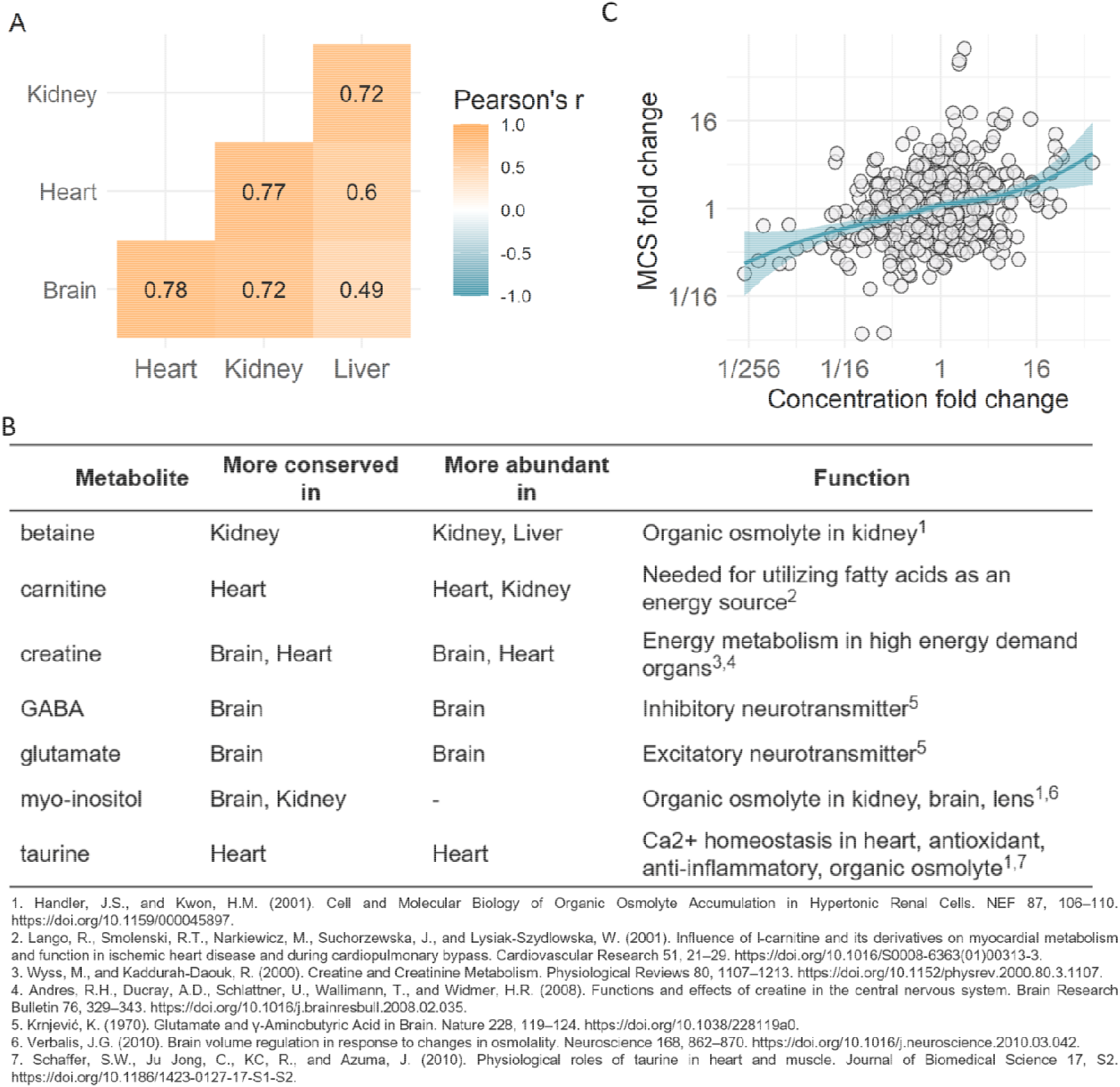
Organ-specific differences in metabolite conservation score. **A**: Similarity of conservation scores of metabolites across the four organs as measured by Pearson’s correlation coefficients. All organ comparisons are highly significant (p< 1.1e^-10^, N=113-122). **B**: Example of shifts in metabolite conservation among organs that likely reflect organ-specific functions. Table shows organ-specific conservation and abundance patterns of selected metabolites (i.e. indicating the organs in which a specific metabolite is more conserved or abundant relative to the cross-organ average) and a description of their organ-specific functions. **C**: Between-organ differences in metabolite concentrations show a positive correlation with between-organ conservation score differences (Spearman’s rho = -0.27, p<10^−4^ from permutation test). For each metabolite, concentration difference (fold-change) and conservation score difference (fold-change) was calculated for all six possible organ pairs. Each dot represents the comparison of two organs for a particular metabolite. Line indicates LOESS regression. Statistical significance was assessed by permutation (Methods).

Several metabolites display elevated concentrations in a particular organ, independent of the species^6^. Given that abundant metabolites tend to show enhanced conservation within organs (see Fig. 2D), we hypothesize that a particular metabolite should be more conserved in organs where it is more abundant. Indeed, several metabolites displaying organ-specific conservation also have higher concentrations in those organs where they are more conserved (Fig. 3B). As a systematic test, we examined the relationship between the relative differences (fold-change) of metabolite concentrations and conservation scores among different organs (Fig. 3C). As expected, between-organ concentration differences and between-organ conservation score differences display a significant positive correlation (Fig. 3C). Note that since the total amount of evolutionary changes in the metabolome varies by organ^6^, we compared the conservation of individual metabolites across organs while accounting for this metabolome-wide effect (Methods).

Together, these results indicate that metabolites vary in their conservation due to differing functional constraints and highlight the key influence of abundance on metabolite conservation.

### Systems modelling illuminates the mechanism of functional constraint

Why are the metabolites that are abundant or involved in essential reactions highly conserved? Metabolite concentrations are principal determinants of reaction rates (fluxes) in the network^3^. As metabolic fluxes obey optimality principles^24^, we propose that selection to maintain key metabolic fluxes at optimal values constrains the evolution of metabolite concentrations, and may explain the higher conservation of metabolites that are abundant or involved in essential reactions. To test this, we simulated evolution in a physiologically relevant mathematical model of central metabolism. We employed a kinetic model of the core metabolism of human erythrocytes, which includes glycolysis, the 2,3-bisphosphoglycerate shunt and the pentose-phosphate cycle, with 40 internal metabolites^25^. The model allowed us to simulate the effects of changes in enzyme kinetic parameters (*i*.*e*. mutations) on steady-state fluxes and metabolite concentrations (Methods). Assuming that the non-mutated model represents a fitness maximum resulting from past evolutionary optimization of erythrocyte metabolism, we approximated the deleterious effect of mutations by calculating deviations in four specific fluxes, referred to as *key fluxes*, that are important for proper erythrocyte functioning^25^ (Methods). Then we simulated evolution with and without stabilizing selection on these key fluxes, the latter corresponding to evolution under pure genetic drift, using a Markov chain Monte Carlo approach^26^ (Suppl. Fig. S9, Methods). Crucially, these evolutionary scenarios do not invoke adaptation towards new optimal states, and hence represent non-adaptive modes of evolution.

As expected, *in silico* metabolite conservation scores are much higher in the presence of stabilizing selection (Suppl. Fig. S10), demonstrating that many metabolome-altering mutations are harmful. Furthermore, between-metabolite differences in MCS increase significantly under stabilizing selection (Suppl. Fig. S10), indicating that the requirement to maintain key fluxes imposes varying levels of constraint across different metabolites. Remarkably, metabolite abundance and involvement in essential reactions are determinants of *in silico* MCS. In particular, we found a significant positive correlation between a metabolite’s abundance in the wild-type model and its conservation score in the simulations (Fig. 4A). Similarly, metabolites involved in reactions that are deemed essential *in silico* (*i*.*e*. reactions that have a large impact on key fluxes when inactivated) show high conservation scores in the simulations (Fig. 4B, Methods). Importantly, these associations hold only in the presence of stabilizing selection, indicating that they are not caused by mutational variability (Fig. 4A-B). Furthermore, metabolite abundance and reaction essentiality are independently associated with *in silico* MCS (Suppl. Table S7).

**Figure 4:**
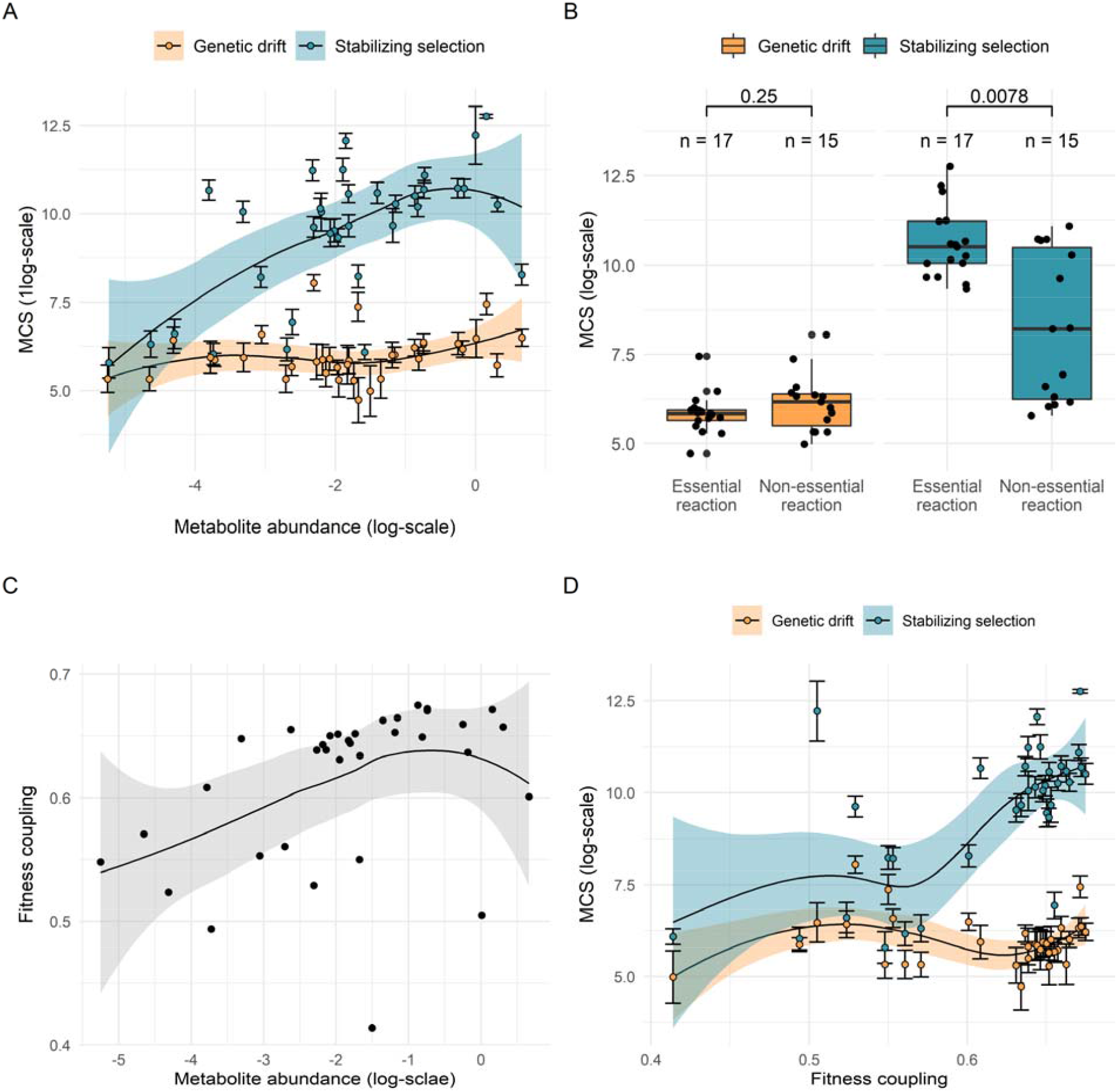
Functional constraints in an in silico model of metabolic evolution. **A:** Metabolites with a higher abundance in the wild-type model show higher conservation scores in the presence of stabilizing selection (blue dots, Spearman rho = 0.55, p = 6.63e-4, N = 35), but not in the absence of selection (orange dots, Spearman rho = 0.30, p = 0.078, N = 35). Each dot and error bar represents the mean and standard deviation of the metabolite conservation score, calculated for a particular metabolite based on 10 simulations. **B:** Metabolites involved in essential reactions (i.e. their products or substrates) have higher conservation scores than those involved in non-essential reactions in the presence of stabilizing selection (blue), but not in the absence of selection (orange), as indicated by two-sided Wilcoxon tests (p = 7.82e-3 and p = 0.246, respectively). Each dot represents the mean conservation score for a particular metabolite based on 10 simulations. **C:** Wild-type abundance of metabolites correlates with their extent of fitness coupling (Spearman rho = 0.49, p = 3.14e-3, N = 35). **D:** Metabolites conservation scores correlate with the extent of fitness coupling under stabilizing selection (Spearman rho = 0.55, p = 7.78e-4, N = 35), but not in the absence of it (Spearman rho = 0.06, p = 0.724, N = 35). The lines in panels A, C and D represent LOESS regressions, with their 95% confidence intervals shown. The boxplots in panel B show the median, first and third quartiles, with the whiskers showing the values within a 1.5 interquartile range distance from the 1^st^ and 3^rd^ quartiles.

We next hypothesized that mutations altering abundant metabolites are more likely to perturb key fluxes and are therefore selected against. To test this, we defined a measure of *fitness coupling* for each metabolite by simulating the impact of single mutations and calculating how strongly changes in the concentration of each metabolite are correlated with changes in key fluxes (Methods). Consistent with the hypothesis, abundant metabolites are indeed more strongly coupled to fitness (Fig. 4C). Furthermore, the extent of fitness coupling of a metabolite correlates with its conservation score inferred under stabilizing selection, but not under pure genetic drift (Fig. 4D). In addition, the four key fluxes assumed to be important for fitness^25^ lead to a significantly stronger correlation between metabolite abundance and fitness coupling than randomly defined key fluxes (Fig. S11A; Methods). Thus, the strong coupling of highly abundant metabolites to fitness is not a by-product of the modelling procedure, but specifically holds for a realistic fitness definition. Further analysis confirmed that the strong conservation of abundant metabolites is mediated by fitness coupling (Fig. S11B).

Finally, the above results also hold for a different model of erythrocytes and a model of human hepatic glucose metabolism (Appendix S1). Overall, these findings indicate that metabolites that are highly abundant or participate in essential reactions are more conserved in their concentrations because they are more crucial to maintain key metabolic fluxes. Importantly, as our evolutionary simulations were devoid of beneficial mutations, non-adaptive evolution is sufficient to explain major patterns of metabolome conservation.

### Evolutionary conservation informs on disease association

Next we asked whether the conservation of a metabolite informs on its involvement in diseases. We first focused on human inborn errors of metabolism (IEMs), which are genetic disorders caused by disruption of specific metabolic pathways^27^. The early onset and high severity of these disorders suggest that metabolites involved in IEMs might be highly constrained in mammals. We compiled metabolites known to be involved in the disease etiology or the diagnosis of IEMs routinely measured in newborn screening (see Methods, Suppl. Table S8). We found that IEM-associated metabolites show significantly higher conservation scores than the rest of metabolites in all four organs (Fig. 5A and Supp. Fig. S12). The strong evolutionary conservation of IEM-associated metabolites is not explained by abundance, a particular class of conserved metabolites or specific metabolic pathways (Suppl. Table S9), suggesting that it reflects their importance for normal metabolic functioning.

**Figure 5:**
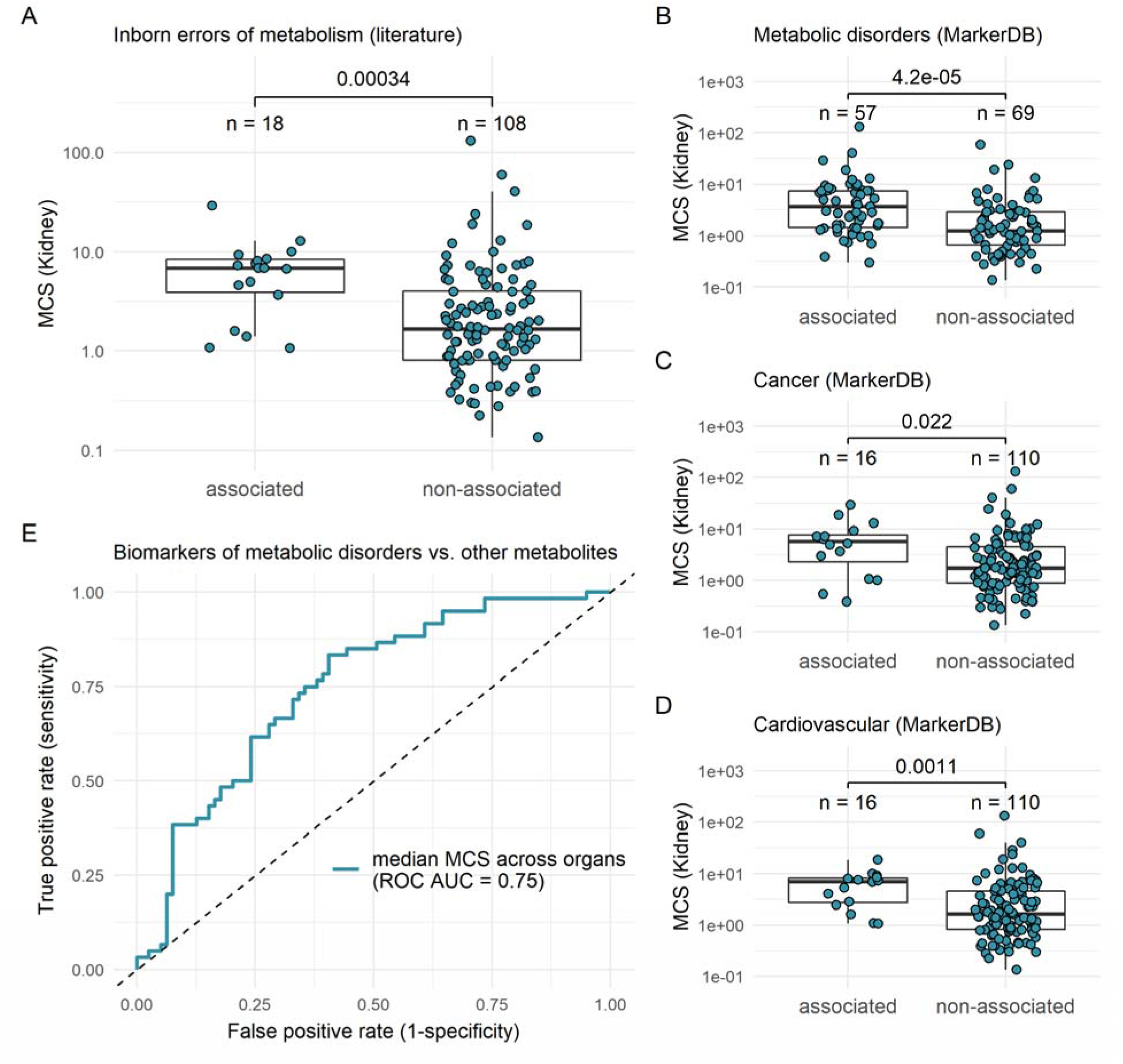
Concentrations of disease associated metabolites are highly conserved. **A:** Metabolites involved in inborn errors of metabolism (IEM) show significantly higher conservation scores than the rest of metabolites in the kidney (two-sided Wilcoxon rank sum test). For other organs, see Suppl. Fig S12. **B-D:** Metabolites associated with three broad diseases conditions (metabolic disorders, cancer, cardiovascular disorders) show a significantly higher level of conservation than the rest of metabolites in kidney (p-values from the Wilcoxon rank sum tests are shown in figure). Boxplots show the median, first and third quartiles, with the whiskers indicating the values within a 1.5 interquartile range distance from the 1^st^ and 3^rd^ quartiles. **E:** Receiver operating characteristic curve (ROC) for prediction of biomarkers of metabolic disorders based on the median conservation score across organs.

To test whether this finding applies to other diseases beyond inborn errors of metabolism, we used MarkerDB, a comprehensive database of clinical biomarkers^28^ and focused on 11 broad disease conditions with sufficient numbers of metabolites (Suppl. Table S10). As expected, metabolites associated with metabolic disorders, including many IEMs, tend to be highly conserved (Fig. 5B). More remarkably, we identified two additional broad disease conditions – cancers and cardiovascular diseases – that are independently associated with highly conserved metabolites (Fig. 5C-D, Suppl. Table S10, Methods). For instance, choline, an essential nutrient and precursor in lipid metabolism, is highly conserved in all four organs, but is not associated with any IEMs (Suppl. Fig. S13). Notably, abnormal choline metabolism is a general hallmark of cancers, and both phosphocholine and total choline-containing metabolite concentrations are used to detect malignant tumors^29^. Furthermore, the oncometabolite succinate^30^ also shows marked conservation in several organs (Suppl. Fig. S13).

Biomarkers of metabolic disorders show the strongest signal of conservation (Fig. 5B), suggesting that evolutionary conservation could potentially be leveraged to identify such biomarkers independent of prior clinical knowledge. As a preliminary test, we computed an aggregate score of conservation across the four organs (Methods). Encouragingly, this conservation score alone separates biomarkers of metabolic disorders from the rest of metabolites with reasonable accuracy (AUC ROC = 0.75, Fig 5E).

Finally, we hypothesized that metabolites involved in multiple diseases are more likely to affect organismal fitness when altered and hence are under stronger stabilizing selection. Indeed, the conservation score shows a positive correlation with the number of specific diseases involving a particular metabolite (Suppl. Fig. S14; effect is independent of abundance and essentiality, Suppl. Table S11).

### Metabolome conservation in the *Drosophila* genus

To test the generality of our main findings, we also analysed metabolome evolution in the distantly related genus *Drosophila*, across a 50-million-year phylogeny (Fig. 6A). We calculated metabolite-specific conservation scores using data on 92 non-lipid metabolites measured in whole adults of 11 *Drosophila* species under the same controlled environment^31^ (Methods). Just as in mammals, MCS varies extensively across different metabolites, spanning over three orders of magnitude (Fig. 6B). Remarkably, using an independent dataset of absolute metabolite concentrations in *D. melanogaster*^32^, we found a significant positive correlation between abundance and the conservation score, with an effect size similar to that in mammals (Fig. 6C). Furthermore, metabolites involved in human IEMs tend be highly conserved in *Drosophila* as well (Fig. 6D). We conclude that metabolome evolution is governed by similar principles in two distant animal phyla.

**Figure 6:**
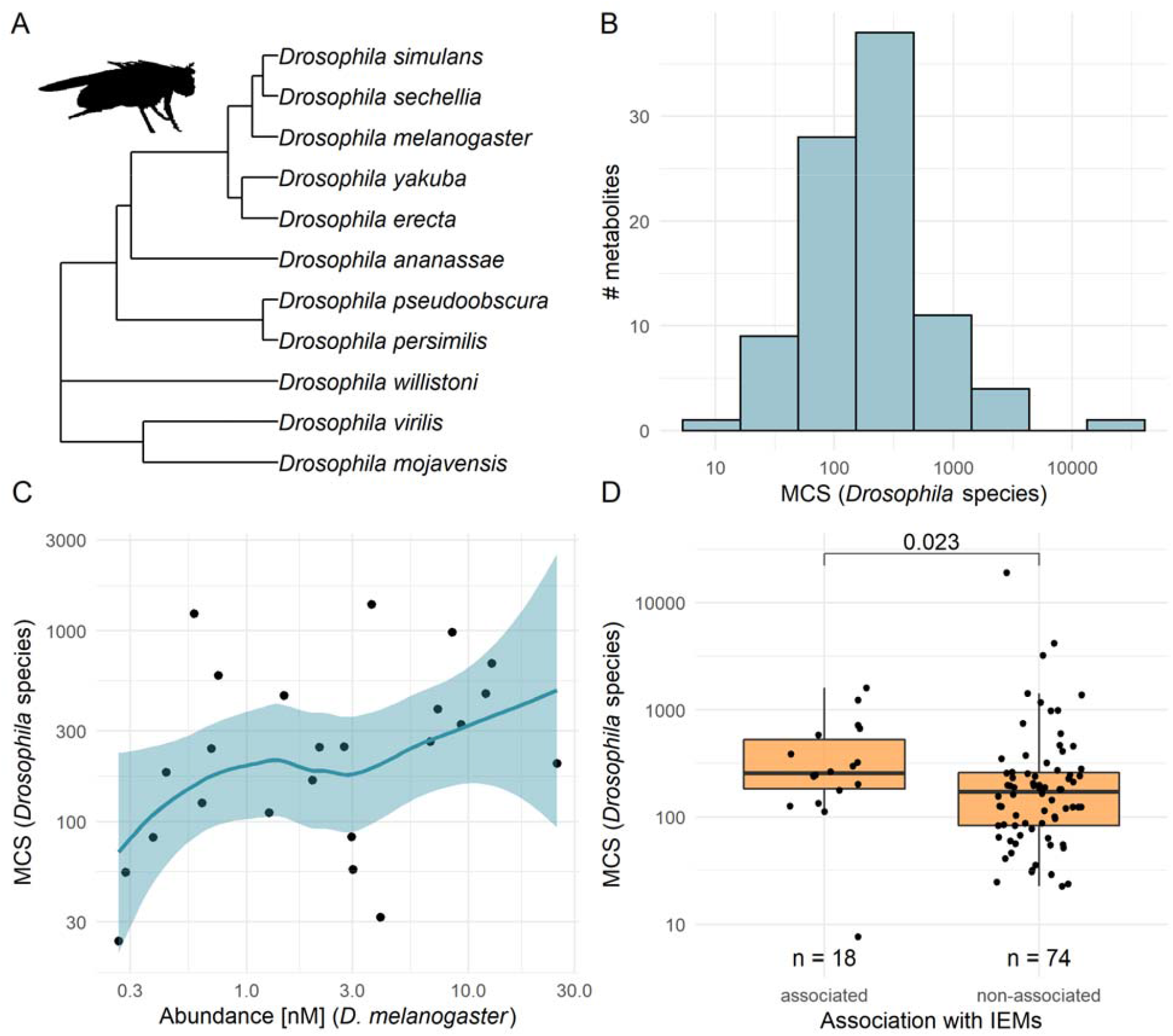
Metabolite concentration conservation in the Drosophila genus. **A**: Phylogenetic tree of 11 Drosophila species included in the analysis, according to^31^. **B**: Distribution of metabolite conservation scores across 92 metabolites, calculated for the Drosophila genus. **C**: Metabolites with a high absolute concentration (abundance) in D. melanogaster tend to show high conservation score in Drosophila (Spearman’s correlation rho = 0.43, p = 0.037, N=24). **D**: Metabolites associated with inborn errors of metabolism in human have an elevated conservation score in Drosophila (two-sided Wilcoxon rank sum test, p-value shown in figure).

## Discussion

In this work, we combine phylogenetic analysis of metabolome data with systems biology modelling to seek general principles governing the evolution of intracellular metabolite concentrations in animals. By introducing a measure of evolutionary conservation of individual metabolite concentrations, we showed that the extent of conservation is largely invariant between closely related clades, but varies extensively across metabolites. Such variation in conservation is predictable based on a few metabolite properties and is consistent with a simple model where natural selection preserves flux through key metabolic reactions while permitting the accumulation of selectively neutral changes in enzyme activities. We further demonstrated that this general conceptual framework of metabolome conservation informs on disease associations and biomarker status of metabolites.

Metabolite abundance emerged as the main determinant of conservation, with highly abundant metabolites displaying the highest level of conservation. Importantly, any particular metabolite displays stronger conservation in organs where it is more abundant, demonstrating that abundance *per se* affects conservation. Systems modelling further showed that abundant metabolites are subject to stronger functional constraints (Fig. 4). Why should it be so? First, the concentrations of highly abundant metabolites might be more rate-limiting for key fluxes than those of low-abundance metabolites, implying a causal effect of metabolite concentration on fitness. This would be analogous to the observation that halving gene dosage is generally more deleterious for highly expressed genes^33^. Alternatively, abundant metabolites might not be particularly rate-limiting, but might be subject to stronger indirect selection due to the harmful side effects of mutations affecting their concentrations^34^. We speculate that there might be less ways to alter abundant metabolites without also perturbing key fluxes, resulting in stronger indirect selection on these molecules. Finally, regardless of their specific functions, abundant metabolites incur broad cellular costs due to limitations on osmotic pressure^11^ and total dry mass^10^, potentially constraining their evolution. Clearly, further studies are needed to test these scenarios.

Our work has profound implications for the neutral theory of molecular evolution, which posits that most within- and between-species variations at the molecular level are selectively neutral rather than adaptive^18^. While the theory explains many aspects of sequence and gene expression evolution^18,35,36^, it has been unclear whether it applies to variations at the metabolome level, which is more closely related to phenotypes^37^. Our results are broadly consistent with a neutral model of metabolome evolution. First, metabolite conservation scores are largely constant across different mammalian clades, suggesting that similar evolutionary forces shape the metabolome despite extensive phenotypic divergence. Second, conservation is determined by the functional properties of metabolites, namely abundance, involvement in essential reactions and association with human diseases. As these metabolite properties likely reflect the level of functional constraints, rather than the amount of adaptive evolution, they support the neutral model. Notably, analogous gene properties – expression level, essentiality and disease association^12^ – determine protein sequence conservation, revealing a striking parallel between the selective constraints driving metabolome and protein evolution. Finally, metabolic modelling demonstrated that stabilizing selection on key fluxes is sufficient to explain the strong conservation of abundant and essential metabolites without the need to invoke adaptation to changing environments. Together, these results suggest that a substantial fraction of metabolome differences among mammals, as well as among *Drosophila* species, are neutral and are permitted rather than favored by selection. It remains to be tested whether further predictions of the neutral model are fulfilled, and whether they also hold for other major taxa. Nevertheless, our simulation study represents an important step towards a theoretical framework of metabolic evolution driven by non-adaptive processes.

The evolutionary history of gene sequences and gene expression levels informs on their disease involvement^15,35^. Our work expands this notion to include a new layer of molecular phenotypes by showing that metabolome conservation is predictive of the disease associations of metabolites. Remarkably, biomarkers can be distinguished from non-biomarker metabolites simply based on the comparison of metabolomes across species, without utilizing any prior clinical knowledge. As expected, metabolite conservation appears to be most informative for metabolic diseases that disrupt basic cellular functions and show an early onset, such as inborn errors of metabolism. More intriguingly, metabolites involved in tumorigenesis are also well conserved, suggesting that cancer avoidance might be an important selective force in wild mammals^38^.

We anticipate that evolutionary metabolomics should have at least two possible applications to aid clinical diagnosis. First, it offers a strategy to identify metabolites whose dysregulation matters the most for human health, and therefore could be involved in disease mechanisms or may be used as biomarkers. Given the plethora of assayed metabolites in metabolomic epidemiology studies^39^, evolutionary conservation may help to prioritize them for biomarker identification and further investigations. Second, it might be possible to infer the range of permissible metabolite concentrations from cross-species data, and use this information to detect pathogenic alterations in individual metabolome profiles^35^. As clinical diagnoses typically rely on the measurement of plasma metabolite levels, it is an important open issue whether the concept of evolutionary conservation could be applied to blood metabolomes, which might be more strongly influenced by environmental effects than tissue metabolomes.

In sum, our findings illustrate how evolutionary comparisons of metabolite concentrations on a network scale can be leveraged to study the functional constraints and pathogenic alterations of cellular metabolism.

## Methods

### Calculating metabolite conservation scores

To study metabolome conservation in mammals, we obtained metabolite concentration measurements of 139 non-lipid metabolites from a multispecies study^6^. The dataset contains relative metabolite concentrations across 26 mammalian species in 4 organs (brain, heart, kidney and liver) and is based on targeted metabolomics measurements involving three distinct liquid chromatography-mass spectrometry (LC-MS) methods. Note that the measured samples were homogenates of freshly frozen tissues of sacrificed animals, matched by age (i.e. young adults) and sex^6^. We calculated the mean of the normalized (log_10_-transformed) relative metabolite concentrations across all biological replicates so that each metabolite in each species and organ is represented by a single relative concentration value. We then used these values as continuous molecular traits for which conservation scores are computed.

To calculate the metabolite conservation scores of individual metabolites, we first fit a simple Brownian Motion model of trait evolution on the concentrations of each metabolite in each organ across the phylogeny, using the fitContinuous function in the Geiger R package. The evolutionary rate parameter of the Brownian Motion model measures the rate of trait diversification along the phylogeny and is in the units of trait variance increase per unit evolutionary time (as approximated by phylogenetic distance). It has been argued that the rate parameter of a simple Brownian motion model is a useful measure of the ‘effective rate’ of trait evolution, even if more complex evolutionary models fit a given trait better^16^. Next, by taking the inverse of the evolutionary rate parameter, we define a measure of metabolite conservation, where metabolites that diverge more slowly in their concentrations over time are represented by higher conservation scores.

Conservation scores were calculated in a similar fashion for relative metabolite concentrations measured in fibroblast cell cultures^17^ and *Drosophila* species^31^ as well. The phylogenetic trees used in these calculations were obtained from^17,31^ and from http://www.timetree.org/^40^.

### Variation of conservation scores among different metabolites versus biological replicate measurements

To assess the impact of measurement noise and / or within-species variation on the inferred metabolite conservation scores and compare it to among-metabolite variation, we made use of multiple biological replicate measurements. Specifically, we sampled randomly with replacement concentration values from 2 to 4 biological replicate measurements depending on the species and the organ, and re-calculated metabolite conservation scores 100 times (*i*.*e*. bootstrap procedure). For each organ, metabolite/species pairs having only 1 replicate were removed from the analysis (89.1% of all organ – species - metabolite triplets have more than 1 replicate). For each organ, we then applied a one-way ANOVA test on the resulting metabolite conservation score distributions to partition the amount of total variance in conservation scores into between-metabolites variance and error variance, the latter capturing variation between biological replicate measurements of the same metabolites.

### Metabolite features associated with metabolite conservation score

Seventeen distinct classes of metabolic features were collected in order to probe their relationship with metabolite conservation score. Information on the regulatory roles of metabolites (enzyme activator, inhibitor and co-factor function) was obtained from^41^ which was compiled from the BRENDA database^42^. Information on the chemical properties of metabolites (chemical class, molecular weight, dissociation constants, water solubility, hydrophobicity) were collected from the HMDB and KEGG databases^43,44^. Metabolite toxicity information, in the form of mouse LD50 values (the concentration of the metabolite that is lethal to 50% of specimens, in mg/kg) was collected from the ChemIDPlus database (https://chem.nlm.nih.gov/chemidplus/). Pathway membership and broad position in the metabolic network (biosynthetic, degradation and energy metabolism) were collected from the KEGG and HumanCyc databases, respectively^43,45^. We note that pathway membership was used only for those metabolic pathways that contained at least 5 metabolites for which we had conservation score calculated, yielding 27 pathways in total. Organ-specific absolute metabolic abundance measurements were obtained for mouse from the Mouse Multiple tissue Metabolome DataBase (MMMDB)^20^. Network degree (the number of reactions a metabolite participates in) was determined using a genome-scale reconstruction of the human metabolic network^46^. Metabolites involved in metabolic reactions that are encoded by essential genes were identified using phenotypic data from mouse knock-out lines. In short, we identified genes whose deletion caused either a lethal phenotype or infertility in the Mouse Genome Database^19^. Next, we defined essential reactions as reactions where the majority (> 50%) of the genes associated with the reaction in the human metabolic network are essential. A metabolite was considered part of the essential set if it participates in at least one essential reaction. For brevity, we refer to such metabolites as ‘essential’ metabolites, even though metabolite essentiality cannot be directly measured.

To probe the relationships between metabolite conservation score and individual metabolite features, we used linear regression modelling. For each individual feature, we fitted a linear model that accounts for both the effect of the given feature and the organ in which the conservation scores were estimated (*i*.*e*. organ membership). This allowed us to assess the general effect of each feature on evolutionary conservation across all four organs simultaneously, while accounting for organ-specific global differences in conservation score. The percentage of variance in metabolite conservation score explained by each feature (as shown in Fig. 2A and Suppl. Table. S1) was calculated by subtracting the R^2^ value of a model containing organ-membership as the only predictor variable from the R^2^ value of the model containing both organ-membership and the feature of interest as predictor variables. Each chemical class and KEGG pathway was evaluated separately.

To identify the main determinants of evolutionary conservation while controlling for covariation between the metabolite features, we performed a multivariate analysis as follows. We fitted an initial linear model that included all features and metabolic pathways that individually had a significant effect on conservation scores (nine metabolic features and seven specific metabolic pathways, as shown in Suppl. Table S4). Next, we used a stepwise feature selection (using the step function in R) to identify the most parsimonious linear model that contains the combination of features that provides the best fit based on the Akaike information criterion (AIC). To quantify the contribution of the individual metabolite features to the most parsimonious model (i.e., independent effect), we fitted simpler models by leaving out single features and calculating the decrease in the adjusted R^2^ value. The portion of variance in MCS explained by the combination of metabolite features was determined by subtracting the independent effect of organ membership from the adjusted R^2^ value of the most parsimonious multivariate model. Note that, in order to minimize the number of features in the multivariate model, we used chemical class as a single multilevel factor, instead of multiple binary features.

### Between-organ differences in metabolite conservation score

For all between-organ analyses, we only included metabolites that were measured in all four tissues (110 metabolites of a total of 139). To compare the conservation of metabolites across organs, we first normalized the conservation scores as two of the four organs, brain and heart, are generally more strongly conserved than the others (i.e. show smaller amounts of total evolutionary divergence across the whole metabolome). Conservation scores were first normalized by log_2_ transformation and then centered on zero for each organ. Then, we calculated the organ-specific deviation in conservation for each metabolite by taking the normalized conservation score from one organ and subtracting the mean normalized scores of the other three organs from it. High conservation deviation for a given organ indicates that the metabolite is more strongly conserved in that particular organ compared to other organs. For each organ, we identified the top 10% most strongly deviating metabolite (Suppl. Table S6).

To test whether between-organ differences in metabolite concentrations are generally associated with shifts in metabolite conservation scores, we first calculated, for each metabolite, the differences in the normalized conservation scores between all organ pairs. Next, between-organ differences in metabolite concentrations were determined by calculating the log_2_ fold change of metabolite levels between all organ pairs for each species and then taking the average of the species-specific fold change values. Thus, each metabolite is described by six conservation scores and six metabolite concentration fold change values, corresponding to all six possible comparisons among the four organs. We then quantified the association between all concentration and conservation score fold change values across all metabolites using the Spearman’s rank correlation coefficient. Because the fold change values associated with a given metabolite are not independent from each other, we calculated the p-value of the correlation using a permutation test as follows. We randomly reassigned the organ memberships of metabolite conservation scores and re-calculated the Spearman’s correlation coefficient across 10,000 permutations, to test whether the observed correlation is significantly higher than expected by chance (i.e. one-sided test).

### Evolutionary simulations in a mechanistic model of central metabolism

#### A kinetic model of the core metabolism of human erythrocytes

We used a publicly available kinetic model of the human erythrocyte central metabolism, including glycolysis, the 2,3-bisphosphoglycerate shunt and the pentose-phosphate cycle^25^. This model contains 40 variable metabolites, 38 kinetic reactions, and 166 kinetic parameters (http://jjj.biochem.sun.ac.za/models/holzhutter/). Four specific fluxes, referred to as *key fluxes*, are assumed to be important for the fitness: **(a)** the formation of 2,3-bisphosphoglycerate (flux *v*_9_), **(b)** ATP utilization (flux *v*_16_), **(c)** glutathione (GSH) oxidation (flux*v*_21_) and **(d)** the synthesis of phosphoribosyl-pyrophosphate (flux*v*_26_).

#### Calculating the fitness effect of mutations in the model

The kinetic model allowed us to simulate the effects of changes in enzyme kinetic parameters (*i*.*e*. mutations) on steady-state fluxes and metabolite concentrations. Mutations are approximated by independent random perturbations to the parameters of the 38 kinetic equations. To simulate a single mutational event, a kinetic parameter *p* is selected at random (uniformly among reactions) and its mutant value *p’*_’_ is derived by multiplying it with a factor drawn from a log_10_-normal distribution of variance *σ_mut_*^2^ (ref.^47^):

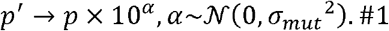

The mutational variance *σ_mut_*^2^ is constant for all the kinetic parameters. The steady-state of each mutant model was computed using Copasi software^48^.

The model’s fitness is approximated by computing the distance *z* between the mutant (MOMA)^49^.The distance *z* represents the deviation from the optimal steady-state in the and wild-type models, similarly to the minimization of metabolic adjustment approach Euclidean space of the relative values of the four key fluxes *v*_9,_ *v*_16,_ *v*_21_ and, *v*_26,_

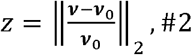

with (*v* = *v*_9,_ *v*_16,_ *v*_21_ *v*_26,_) the key flux levels in the mutant model, σ_0_ the wild-type key flux levels, and where the division by *v*_0_ is element-wise. Importantly, this definition assumes that the wild-type model represents a fitness maximum resulting from past evolutionary optimization of erythrocyte metabolism. Note that such an evolutionary optimization may reflect trade-offs between the maximization of key fluxes, and the minimization of enzymatic production costs and metabolite levels due to molecular crowding, osmotic pressure and other broad cellular costs^10,11^ In this model, any mutation in kinetic parameters is deleterious, as it increases the distance *z*. Moreover, we assume that in the vicinity of the wild-type model, the different kinetic. parameters of the same enzyme can be mutated independently without strongly violating thermodynamic constraints, if mutation sizes are small enough (see parameter values in Suppl. Table S12).

#### Evolutionary simulations using a Markov chain Monte Carlo (MCMC) approach

To simulate evolution, we implemented a Markov chain Monte Carlo (MCMC) modelling algorithm^26^. This approach is assumed to be efficient under the weak mutation-strong selection regime^50^.

As illustrated in Figure S9, starting from the wild-type model (Fig. S9A) and at each iteration *t* of the MCMC algorithm:

1. A kinetic parameter *p* is selected at random and mutated (Eq. 1; Fig. S9B),
2. The steady-state of the mutant model is computed (Fig. S9C). If the mutant does not reach a steady-state, the mutation is discarded
3. The distance *z* between the mutant and the wild-type models is computed (Eq. 2). Stabilizing selection is simulated by applying a selection threshold *w* to the distance *z*. If *z* < ω the mutation is accepted. Else, the mutation is discarded (Fig. S9D). Thus, no mutation can improve the non-mutated model fitness.
4. A new iteration *t* + 1 is computed (Fig. S9E).

We ran 10 repetitions of *T*= 10,000 iterations in two different simulation experiments: **(i) Genetic drift simulations**, where all the mutations are accepted (ω = +∞), and **(ii) Stabilizing selection simulations**, where a selection threshold ω = 1x 10^−4^ is applied on the distance *z* between mutated and non-mutated models. For all the simulations, the mutation size was σ _mut_ =1 ×10^−2^. Simulation parameters are described in Suppl. Table S12.

The numerical framework (as a Python package), simulation results and scripts for additional analyses are publicly available on GitHub (https://github.com/pappb/Liska-et-al-Principles-of-metabolome-conservation

#### In silico metabolite conservation score

At the end of the evolutionary simulation, the evolutionary rate of each metabolite concentration is calculated based on a Brownian motion estimation model^16^,

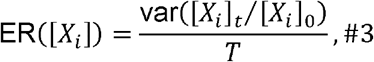

With ER([*X*_*i*_])the evolution rate of the concentration [*X*_*i*_]of metabolite, [*X*_*i*_]_*t*_being its concentration at iteration t,[*X*_*i*_]_0_the concentration of the wild-type model, and *T* the total number of iterations of the simulation. The conservation score of each metabolite is then calculated by taking the inverse of the evolution rate.

#### Calculating the fitness coupling of metabolites

We defined a fitness coupling measure for each metabolite by introducing many independent random single mutations into the kinetic parameters of the model and by calculating the Spearman correlation coefficient between the relative change of metabolite concentrations and the relative change of the four key metabolic fluxes. To this aim, we performed *N* = 10,000 independent single mutations of the wild-type model, by selecting a single kinetic parameter at random uniformly among reactions, and mutating it in a log_10_-normal distribution of size *σ_mut_* ^2^ = 1 ×10^−2^ We measured each time the relative change of metabolite abundances and key fluxes in response to mutations. We then used this result to compute pairwise correlations between fluxes and metabolites. Specifically, for each steady-state flux *v*_j_ and each steady-state metabolite abundance [*X_i_*] across all mutations, the Spearman correlation was computed between the absolute value of relative changes, compared to the wild-type model.

The fitness coupling of a given metabolite X_i_ was then evaluated by computing the mean correlation between the metabolite and the four key fluxes σ= σ_9,_ σ_16,_ σ_21_ and σ_26,_ :

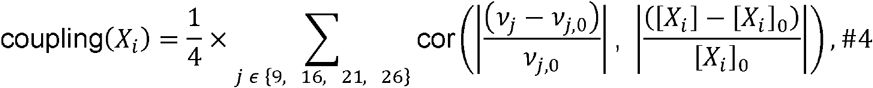

where *v*_*j*,0_ is the wild-type value of *v*_j_, and [*X_i_*]_0_ is the wild-type value of *X_i_*

#### Exploration of random combinations of key metabolic fluxes

We also used the methodology described above to compute the coupling of metabolites (Eq. 4) to 10,000 combinations of key fluxes drawn at random (from random 1-uplets to 4-uplets). For each random combination, the fitness couplings of metabolites were computed, as well as the Spearman correlation between metabolite abundances and their fitness coupling. For 100 random combinations of key fluxes, we also computed one stabilizing selection simulation (*ω* =1 ×10^−4^, *σ_mut_* = 1 × 10^−2^, *T* = 10,000) per combination, in order to compute the Spearman correlation between metabolite abundances and their conservation scores.

#### Calculating in silico reaction essentiality

For each of the 38 reactions of the model, we reduced the flux level to a small fraction of the wild-type level (0.001% for the erythrocyte model from^25^), computed the new steady-state and evaluated the deviation of the four key fluxes relative to their wild-type level. This measure quantifies the “essentiality” of each reaction regarding deviation from optimal key flux levels and hence fitness. We considered a reaction as essential if at least one key metabolic flux is dropped to zero upon its inhibition. Metabolites that are substrates or products of at least one essential reaction are classified as ‘essential metabolites’. We were able to determinate the essentiality of 32 metabolites. It was not possible to calculate it for 3 metabolites because of numerical stability issues.

#### Removal of low-varying metabolites

From all modeling analyses, we excluded five metabolites whose variability was either zero or underestimated in evolutionary simulations: NAD, P1NADPH, glutathione, pyruvate and lactate. These metabolites are insensitive to mutations, as the variabilities of NAD, P1NADPH and glutathione are zero in some genetic drift simulations, while pyruvate and lactate are directly dependent on constant input/output metabolites through transport reactions.

#### Metabolites associated with human diseases

Metabolites associated with inborn errors of metabolism (IEM) were compiled as follows. We included 24 IEM diseases from the US Health Resources and Services Administration’s core recommended uniform newborn screening panel (https://www.hrsa.gov/advisory-committees/heritable-disorders/rusp). We identified disease associated metabolites by manual curation from the relevant literature, as well as the Online Mendelian Inheritance in Man (OMIM) database (https://www.omim.org/) and the Orphanet database of rare diseases (https://www.orpha.net). Any metabolite whose concentration is known to be affected by the disease-causing mutation or is known to show an altered concentration on diagnostic panels was classified as being associated with the IEM disease.

We tested the difference in conservation scores between IEM associated and non-associated metabolites in all four measured organs using ANOVA. To test whether the results are not biased by amino acids, which are prevalent among IEM associated metabolites, we repeated the test after excluding all metabolites that are classified as “Amino acids, peptides, and analogues” according to^6^. To ensure that the high conservation of IEM associated metabolites is not driven by single specific metabolite pathways, we identified five metabolic pathways in KEGG that include three or more IEM associated metabolites: “Alanine, aspartate and glutamate metabolism”, “Arginine biosynthesis”, “Phenylalanine metabolism”, “Valine, leucine and isoleucine biosynthesis” and “Valine, leucine and isoleucine degradation”. We then repeated the ANOVA test 5 times, excluding each one of the above pathways in turn.

For the expanded disease association analysis, we collected chemical biomarkers from the MarkerDB database^28^. The database includes the known chemical biomarkers of a total of 407 human diseases, all of which belong to at least one of 20 broad disease conditions present in MarkerDB. Note that all conditions that are listed in the categories “Others” (such as pregnancy) and “Exposure” (such as smoking) only were omitted from further analysis, as most of these are not strictly disease conditions. In total, 106 metabolites in our dataset were associated with at least one broad disease condition.

To probe the associations between conservation score and involvement of metabolites in broad disease conditions, we focused on 11 broad disease conditions, each of which is associated with at least 10 metabolites in our dataset. The 11 broad conditions include cancers, cardiovascular system disorders, digestive system disorders, endocrine disorders, germ line disorders, haematological- and lymphatic disorders, immune disorders, mental- and behavioural disorders, metabolic disorders, nervous system disorders and urinary system disorders. Because the same metabolite might be involved in multiple broad disease conditions, we used a multivariate approach to determine which disease conditions are significantly associated with metabolite conservation score while controlling for the effects of other disease conditions. First, we determined which broad disease conditions’ biomarkers are significantly more conserved than non-biomarker metabolites using univariate two-sided Wilcoxon rank-sum tests (p<0.05 in at least three out of the four organs). Then, we determined which of the remaining disease conditions show significant independent associations with conservation score using a multivariate linear regression model.

To estimate the extent to which metabolites associated with metabolic disorders in MarkerDB can be predicted based on metabolite conservation score, we first calculated an aggregate conservation score for each metabolite that represents its level of conservation across the four organs. This was achieved by first normalizing the conservation scores in each organ (see *Between-organ differences in metabolite conservation score*), and then taking the median value across the four organs as an aggregate MCS. We then built a classification model using logistic regression that predicts association with metabolic disorders using only the aggregate MCSs. We then evaluated the prediction accuracy of the classifier by a receiver operating characteristics (ROC) curve analysis and by calculating area under the curve (AUC) using the R package “ROCR”^51^.To test the relationship between a metabolite’s conservation score and the number of associated diseases, we used a Spearman’s correlation. This analysis included all specific metabolite - disease associations from MarkerDB, not just those involving the 11 broad disease condition categories.

#### Metabolome conservation in the *Drosophila* genus

Metabolomics data of 92 non-lipid metabolites measured in 11 *Drosophila* species (*D. ananassae, D. yakuba, D. erecta, D. melanogaster, D. simulans, D. sechellia, D. pseudoobscura, D. persimilis, D. willistoni, D. virilis, D. mojavensis*) and the phylogenetic tree describing the evolutionary relationship between the species were obtained from^31^. Conservation scores were computed as described above (see “Calculating the conservation scores of metabolites”)^31^.

Absolute metabolite concentrations of 35 non-lipid metabolites were obtained from^32^. We used the metabolite concentrations measured in whole *Drosophila melanogaster* larvae samples, in order to best match the samples used in^31^. 24 of these 35 metabolites overlapped between the datasets.

## Supporting Information

**Appendix S1**: Evaluation of two additional kinetic models.

**Data S1:** Conservation scores, metabolite features, pathway – and disease biomarker associations based on MarkerDB for 139 metabolites.

## Acknowledgements

We thank Csaba Pál, Martin J. Lercher and László G. Nagy for discussions and critical comments on the manuscript. This work was supported by the National Research, Development and Innovation Office Élvonal Program KKP 129814 (BP), the “Lendület” program of the Hungarian Academy of Sciences LP2009-013/2012 (BP), the European Union’s Horizon 2020 research and innovation program Grant No. 739593 (BP), The Hungarian Academy of Sciences Premium Postdoctoral Research Program (PREMIUM-2018-294 to B.S.), János Bolyai Research Fellowship from the Hungarian Academy of Sciences (BO/00728/21/8 to B.S.), New National Excellence Program of the Ministry of Human Capacities (Bolyai+, UNKP-21-5-SZTE-564 to B.S.), National Research, Development and Innovation Office (PD 128271 to R.T.) and the Tobacco-Related Disease Research Program of the University of California (TRDRP T31FT1772 to G.B.).

## Author information

These authors contributed equally: Orsolya Liska, Gábor Boross, Charles Rocabert

### Author contributions

O.L. and G.B. performed most data analyses with support from C.R. and B.S.. C.R. planned and perfomed the simulation analyses. O.L., G.B., C.R. interpreted results with the assistance of B.S. and R.T.. B.P., O.L., C.R. and G.B. wrote the manuscript with input from all authors. B.P. and G.B. conceived and supervised the project.

### Corresponding authors: Balázs Papp and Gábor Boross

## Competing interests

The authors declare no competing interests.

## Data and materials availability

All data associated with this study are available in the supplementary materials. Data and code associated with the systems modeling work is available on GitHub (https://github.com/pappb/Liska-et-al-Principles-of-metabolome-conservation).

## Notes

https://github.com/pappb/Liska-et-al-Principles-of-metabolome-conservation

### Competing Interest Statement

The authors have declared no competing interest.

