## Supplementary Material for "Principles of metabolome conservation in animals"

**Supplementary files - Principles of metabolome conservation in animals (Liska et al.)**

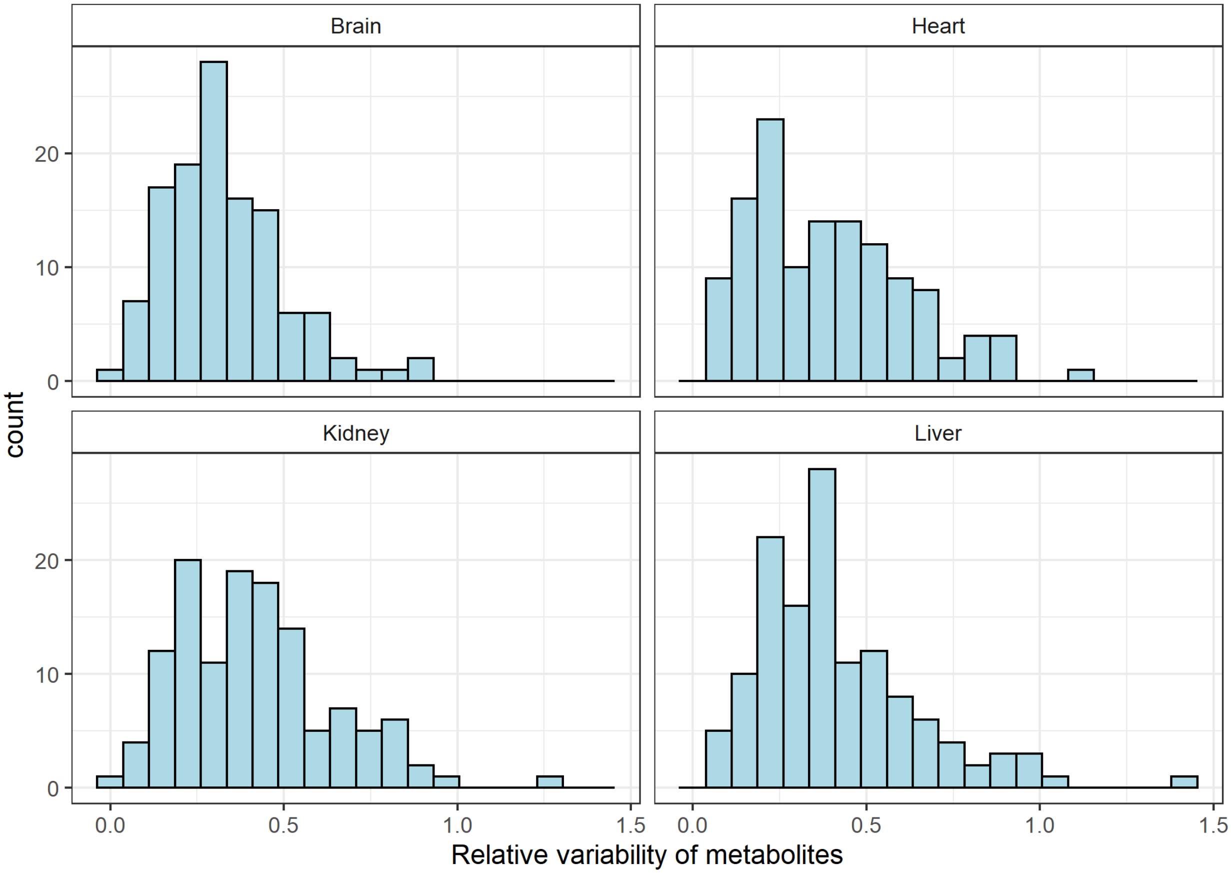


**Figure S1**: **Metabolites differ extensively in their between-species concentration variability.** For each organ, the plot shows the distribution of the relative variability of the concentrations of the studied metabolites calculated across 26 species ($N_{Brain}$=121, $N_{Heart}$=126, $N_{Kidney}$=126, $N_{Liver}$=132 metabolites). Relative variability is defined as the standard deviation of the log10 transformed (normalized) metabolite intensities/measurement[1]. The relative variabilities display up to 23-40-fold differences between metabolites in the four organs (29, 23, 40 and 24-fold for brain, heart, kidney and liver, respectively).

**A** Brain (100 repetitions)

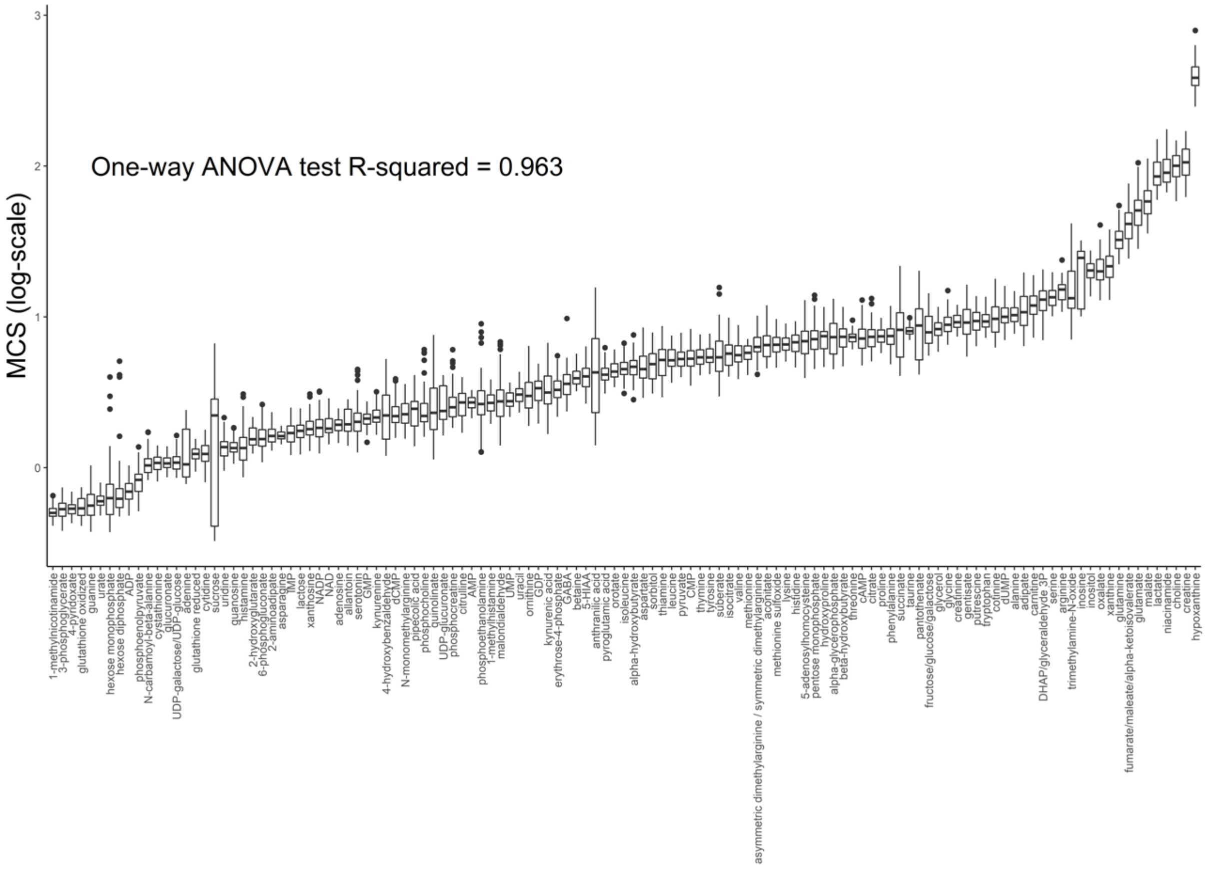


**B** Heart (100 repetitions)

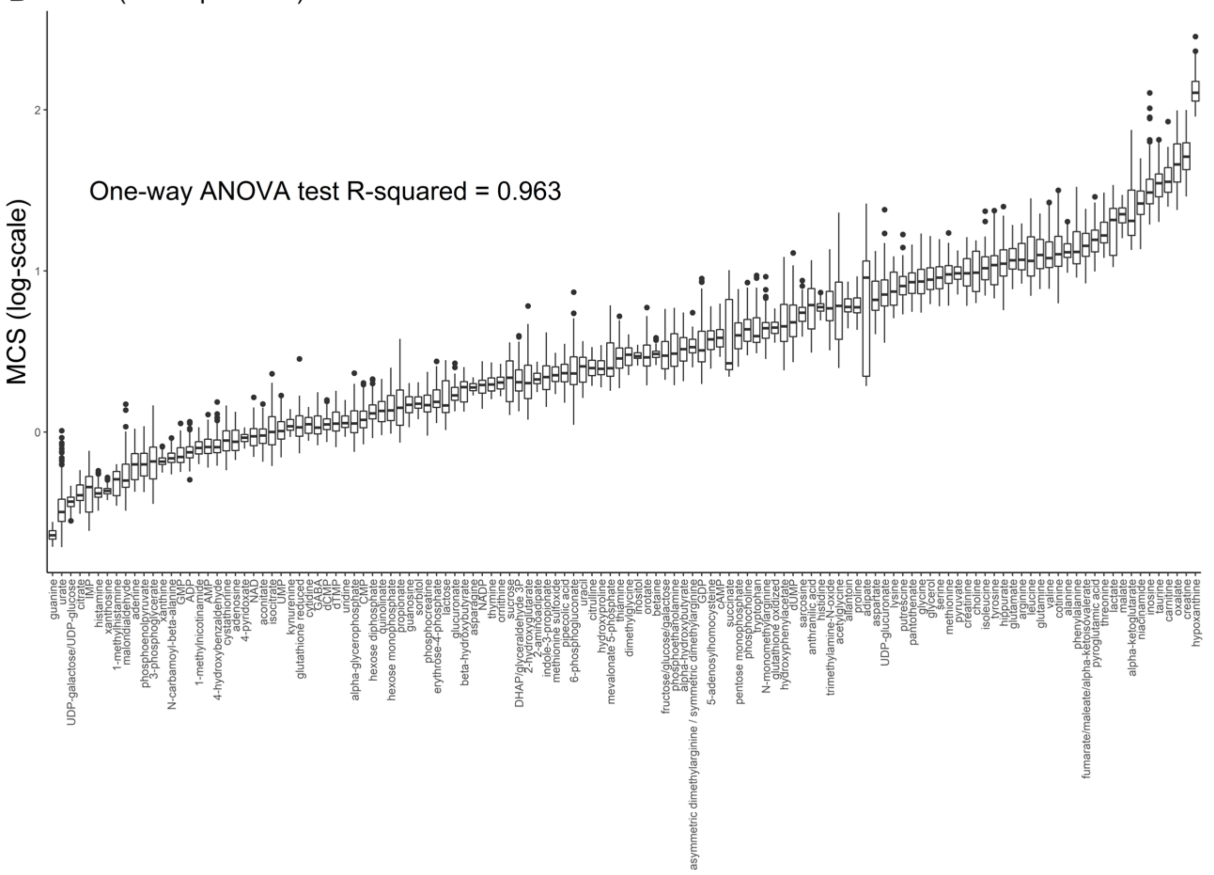

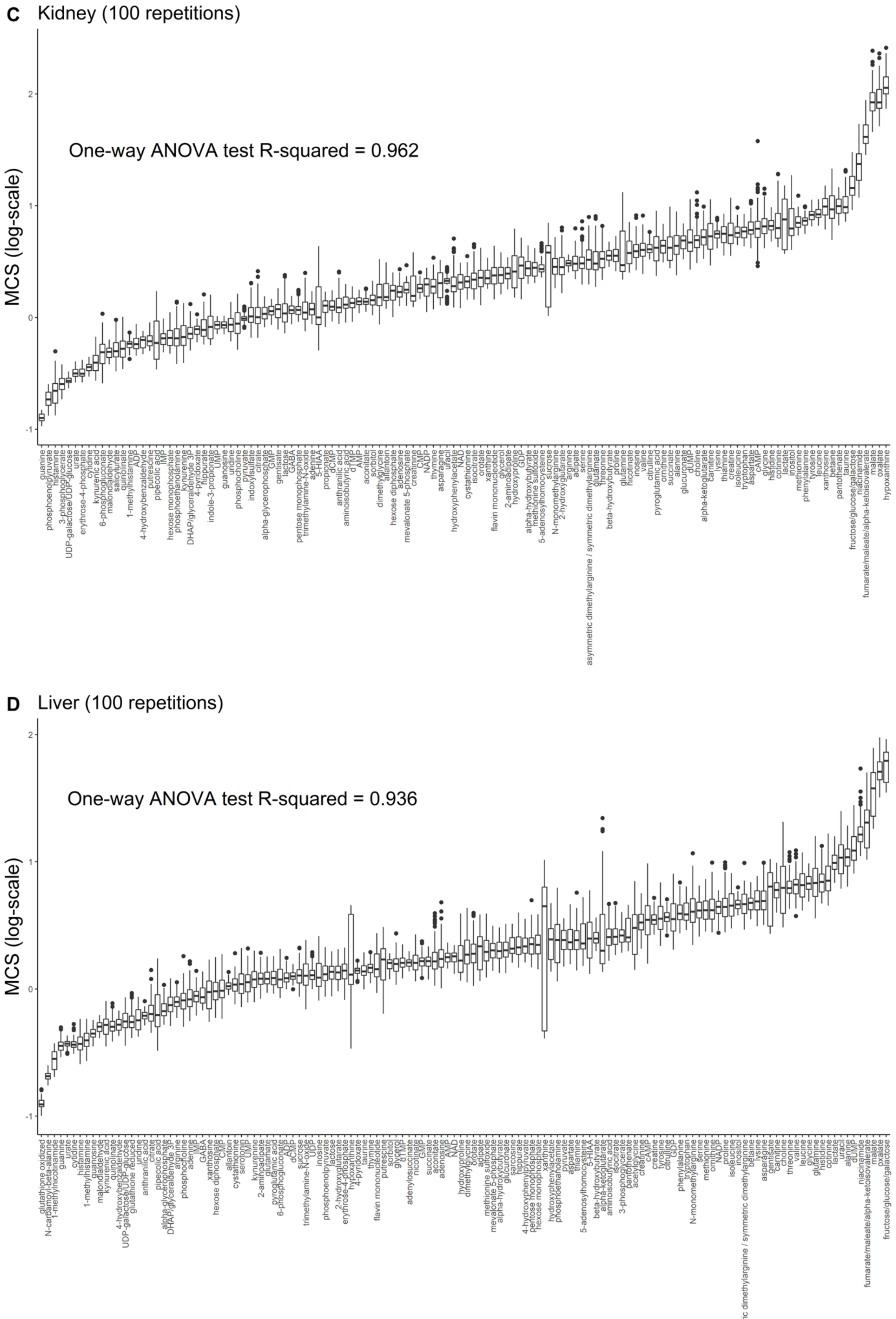

C
Kidney (100 repetitions)
One-way ANOVA test R-squared = 0.962
MCS (log-scale)
D
Liver (100 repetitions)
One-way ANOVA test R-squared = 0.936
MCS (log-scale)

**Figure S2: Variation of MCSs among metabolites versus biological replicate measurements.** For each organ, we computed a one-way ANOVA to estimate the percentage of total variance of metabolite conservation scores (MCS) explained by between-metabolite differences versus differences between biological replicate measurements based on resampling of replicates (see Methods). Box-plots show the distribution of metabolite conservation score estimates calculated from 100 bootstrap samples of biological replicate measurements (see Methods). For each organ, metabolites are ranked by the mean value of their conservation scores. The R-squared values are shown, and all the p-values associated to ANOVA tests are $< 2.2 \times 10^{-16}$. **A** Brain ($R^2 = 0.963$). **B.** Heart ($R^2 = 0.963$). **C.** Kidney ($R^2 = 0.962$). **D.** Liver ($R^2 = 0.936$).

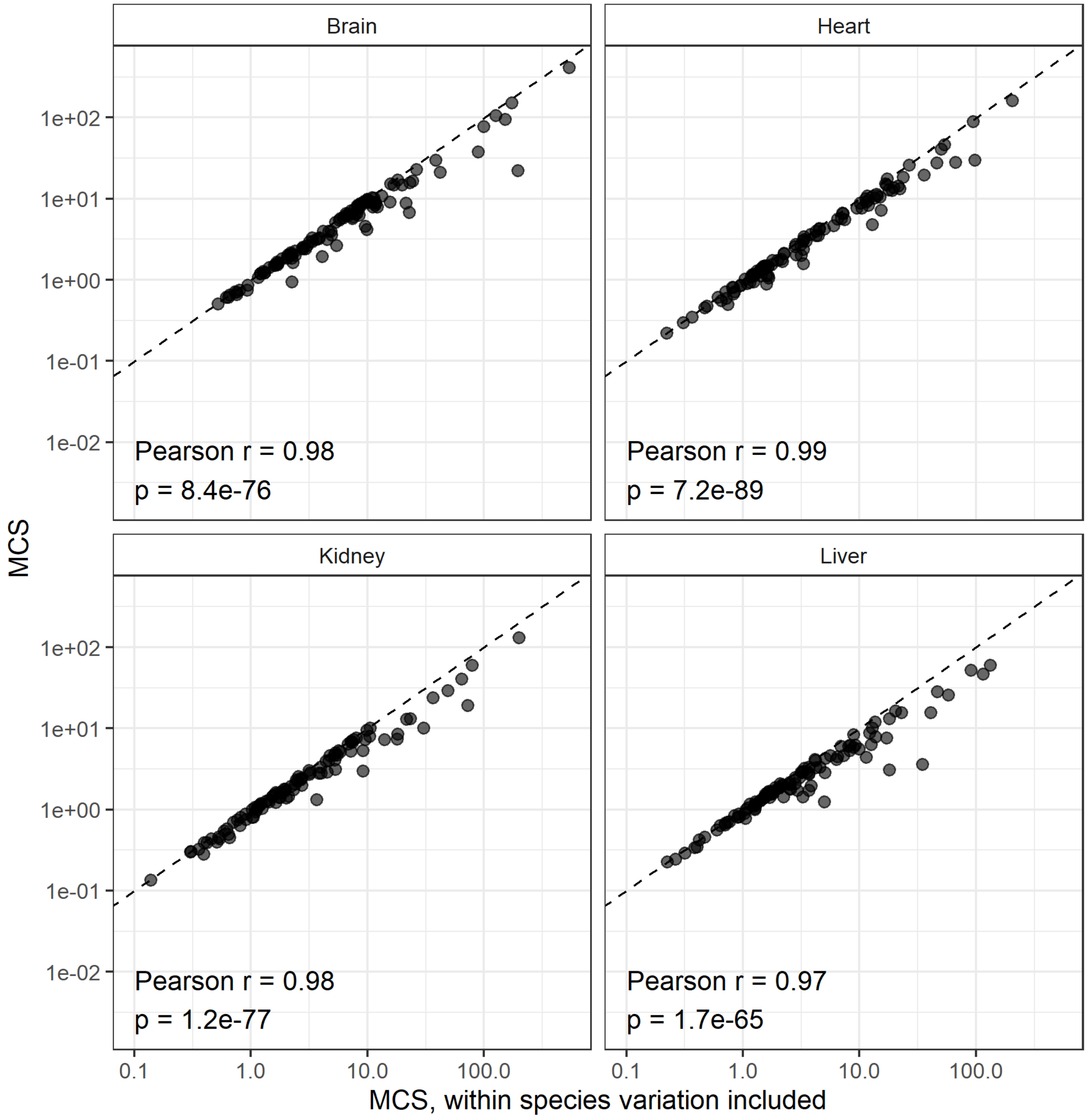


**Figure S3: Conservation score estimates show remarkably strong correlation with those calculated while incorporating within-species measurement noise**. While metabolite conservation scores that account for within-species variation as well are generally lower, they show remarkably strong correlation with the scores calculated based on the metabolite measurements only ($N_{Brain}$ = 113, $N_{Heart}$ = 105, $N_{Kidney}$ = 102, $N_{Liver}$ = 108).

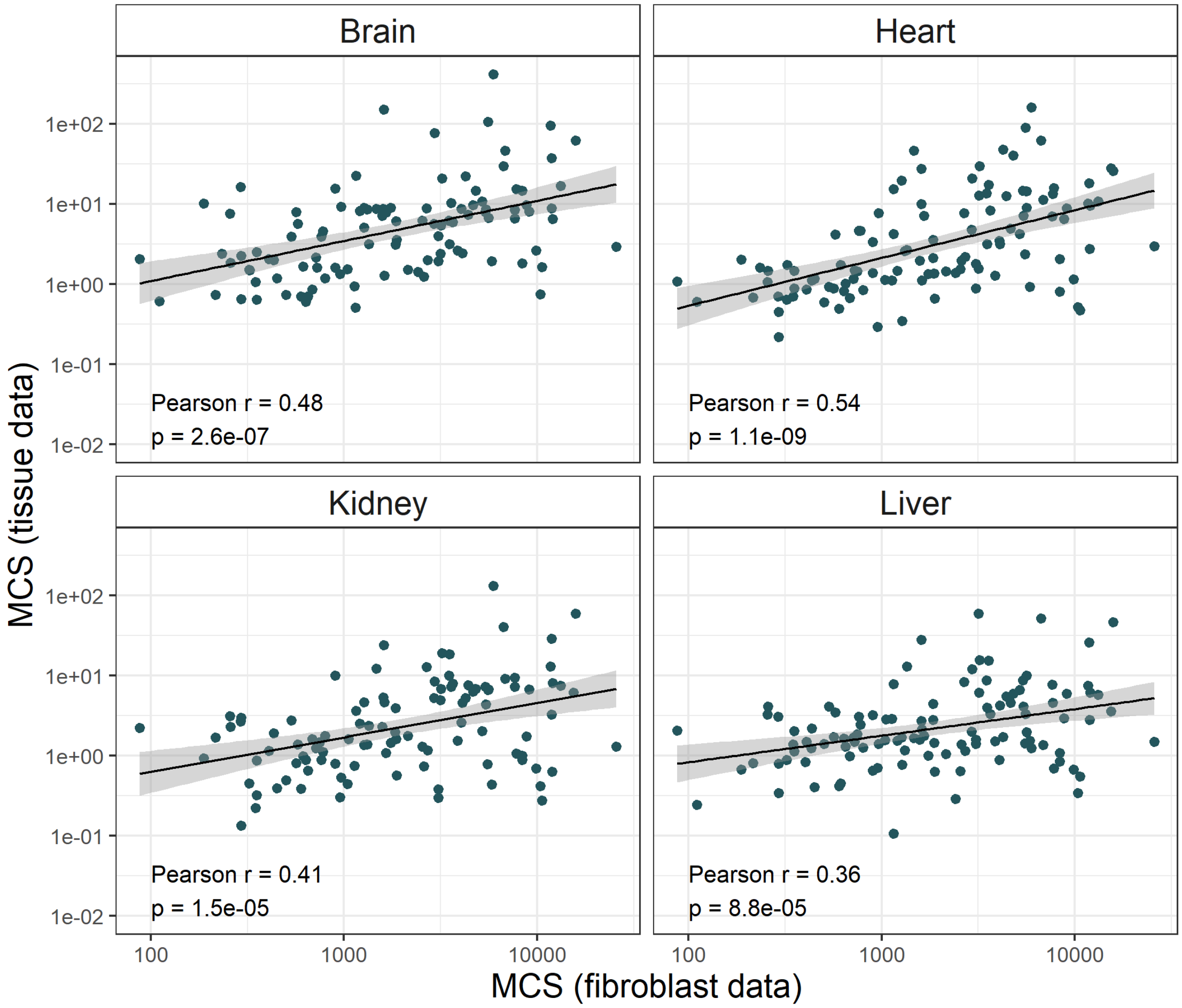


**Figure S4: Metabolite conservation scores inferred from *in vivo* mammalian organ samples correlate with those based on metabolome measurements in cell cultures.** Scatterplots show the correlation between the conservation scores of metabolites in mammalian organ samples and those measured in fibroblast cell cultures ($N_{Brain}$ = 105, $N_{Heart}$ = 110, $N_{Kidney}$ = 106, $N_{Liver}$ = 111). Note that the same metabolomic pipeline was used for both datasets. The lines represent linear regressions, with their 95% confidence intervals shaded in grey. All four organs showed significant positive correlation with fibroblast derived conservation scores.

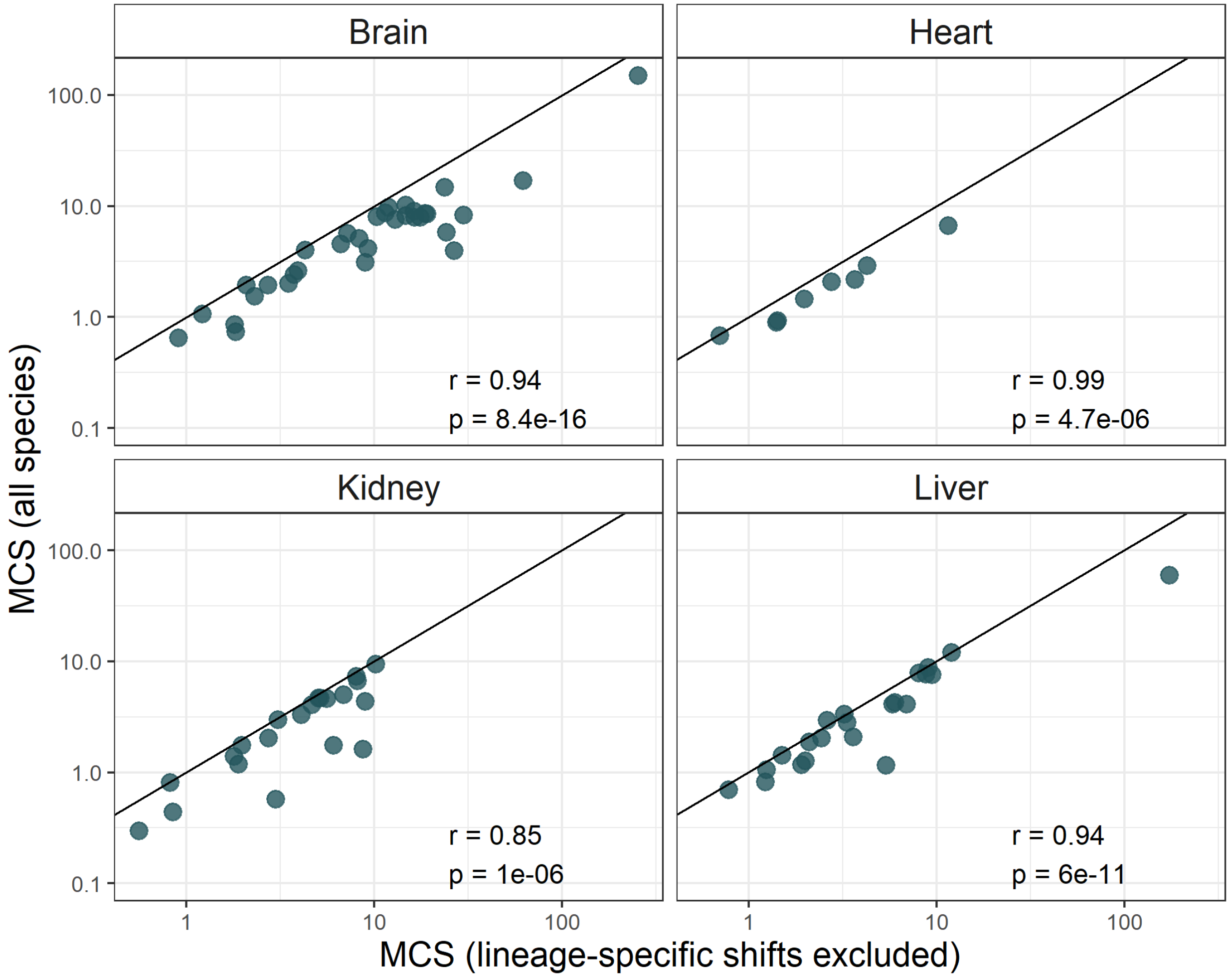


**Figure S5**: **Impact of lineage-specific evolutionary shifts in metabolite concentrations on MCS inferences.** For each organ, we focused on metabolites that show evidence for lineage-specific shifts in concentrations according to[2]. A total of 33, 8, 21 and 22 metabolites show lineage-specific concentration changes in brain, heart, kidney and liver, respectively. For these metabolites, we calculated metabolite conservation scores after excluding those species where lineage-specific selection occurred[2]. In all four organs, these MCSs show strong positive correlations with those calculated based on all 26 species based on Pearson correlations of log-scaled MCSs. The line in the figures depicts the identity function (slope of 1).

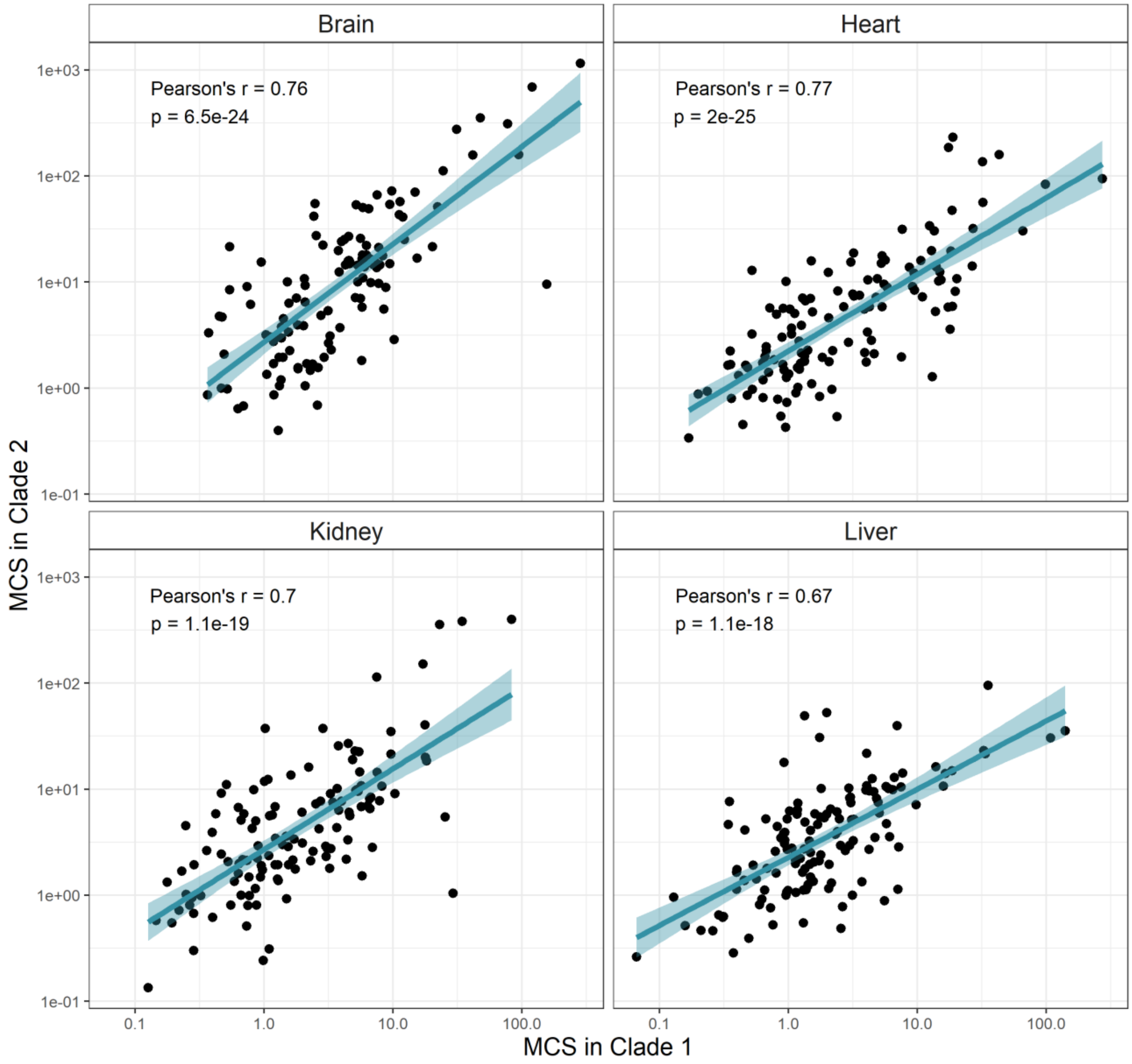


**Figure S6: Metabolite conservation scores calculated on two independent clades of the tree are well correlated in all four organs.** Pearson correlations are significant in all four organs ($N_{Brain}$ = 121, $N_{Heart}$ = 125, $N_{Kidney}$ = 126, $N_{Liver}$ = 132). Lines depict the fitted linear regression. See main figure Fig. 1C for the definition of Clade 1 and Clade 2.

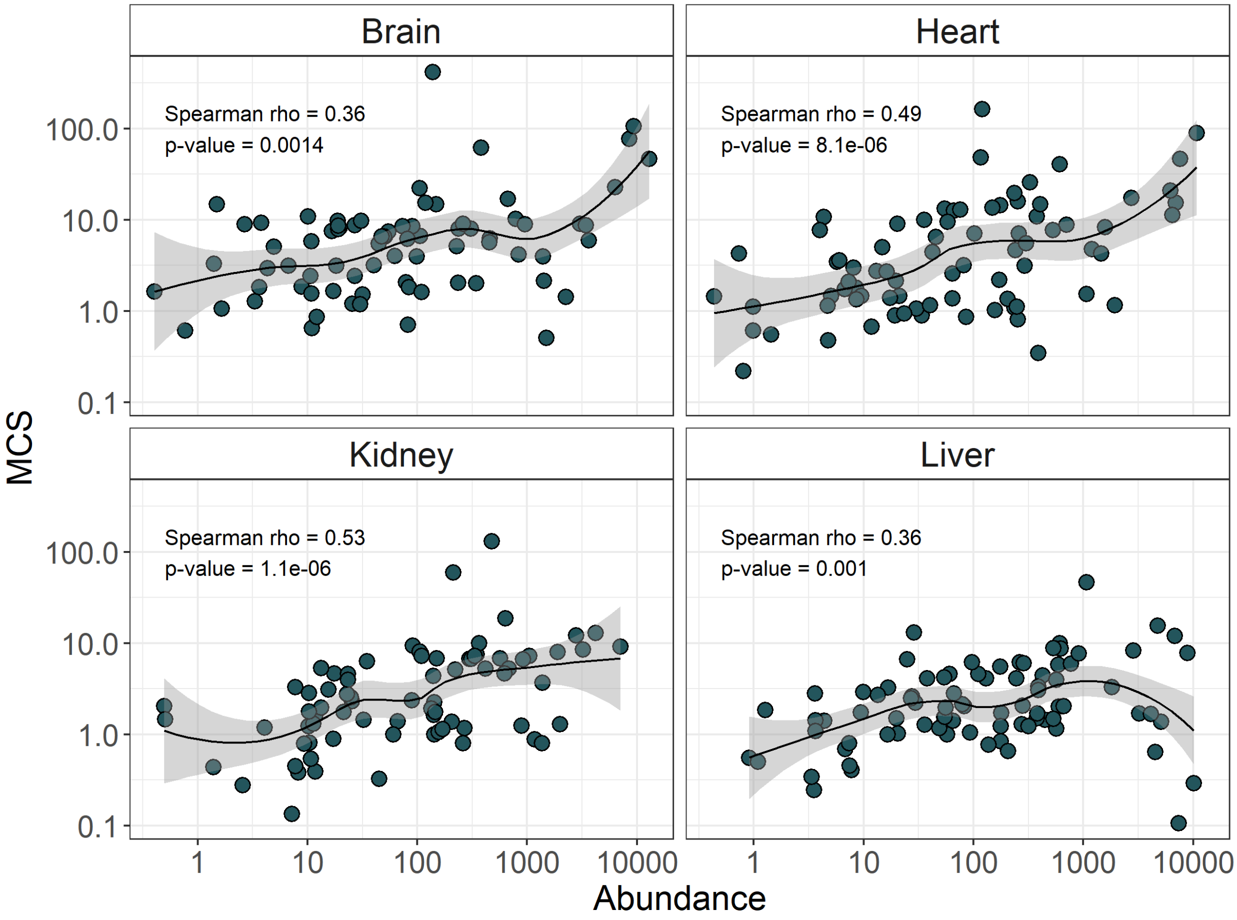


**Figure S7**: **Highly abundant metabolites tend to be more conserved in their concentrations in all organs.** Scatter plots show relationships between metabolite conservation scores of and absolute metabolite abundances (both $log_{10}$-scaled, $N_{Brain}$ = 77, $N_{Heart}$ = 78 $N_{Kidney}$ = 76, $N_{Liver}$ = 83). We used abundance data from an independent study quantifying absolute metabolite concentrations in mice from matched organs[3]. Lines indicate smooth curves fitted using LOESS regression, with their 95% confidence intervals shaded in grey.

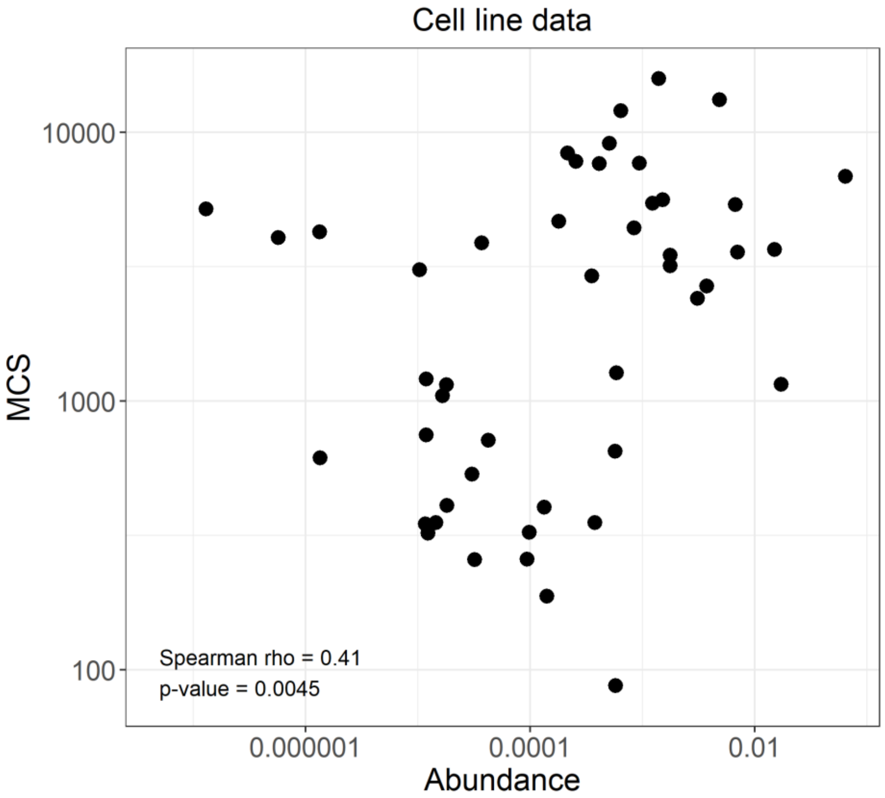


**Figure S8**: **Cell culture-based metabolome data confirms the high MCS of abundant metabolites**. Metabolite conservation scores were calculated from metabolome measurements of primary skin fibroblasts of 16 mammalian species (relative concentrations)[4]. As absolute metabolite concentration was not available for fibroblast cultures, we used absolute concentration data on a mouse cell line derived from baby mouse kidney cells (iBMK)[5]. We found a significant positive correlation between abundance and conservation (Spearman's rho = 0.41, p-value = 0.0045, N = 47).

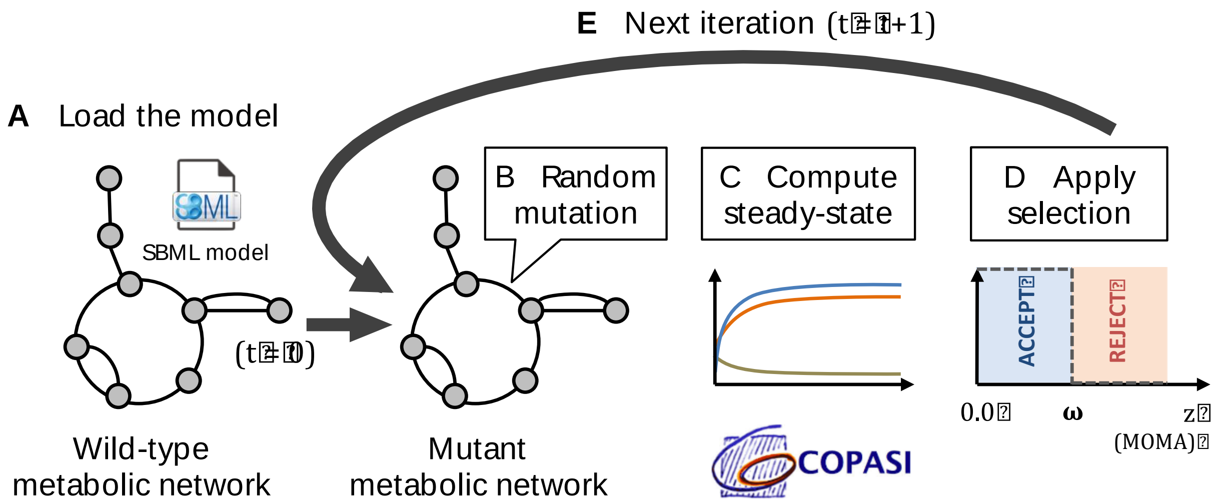


**Figure S9: Overview of the algorithm to simulate the evolution of metabolite concentrations. A.** The model of interest is loaded as a wild-type from an SBML file[6]. Kinetic equations, kinetic parameter values and initial metabolite concentrations must be specified. **B.** At each iteration $t$, a single kinetic parameter is selected at random and mutated through a $\log_{10}$-normal distribution of standard deviation $\sigma_{mut}$. **C.** The new steady-state is computed using the Copasi software[7], and the deviation $z$ of key metabolic fluxes in the mutant relative to their wild-type is computed (see Methods). **D.** If $z$ is under a given selection threshold $\omega$ ($z < \omega$) the mutation is accepted. Otherwise, the mutation is rejected. **E.** A new iteration $t + 1$ is computed.

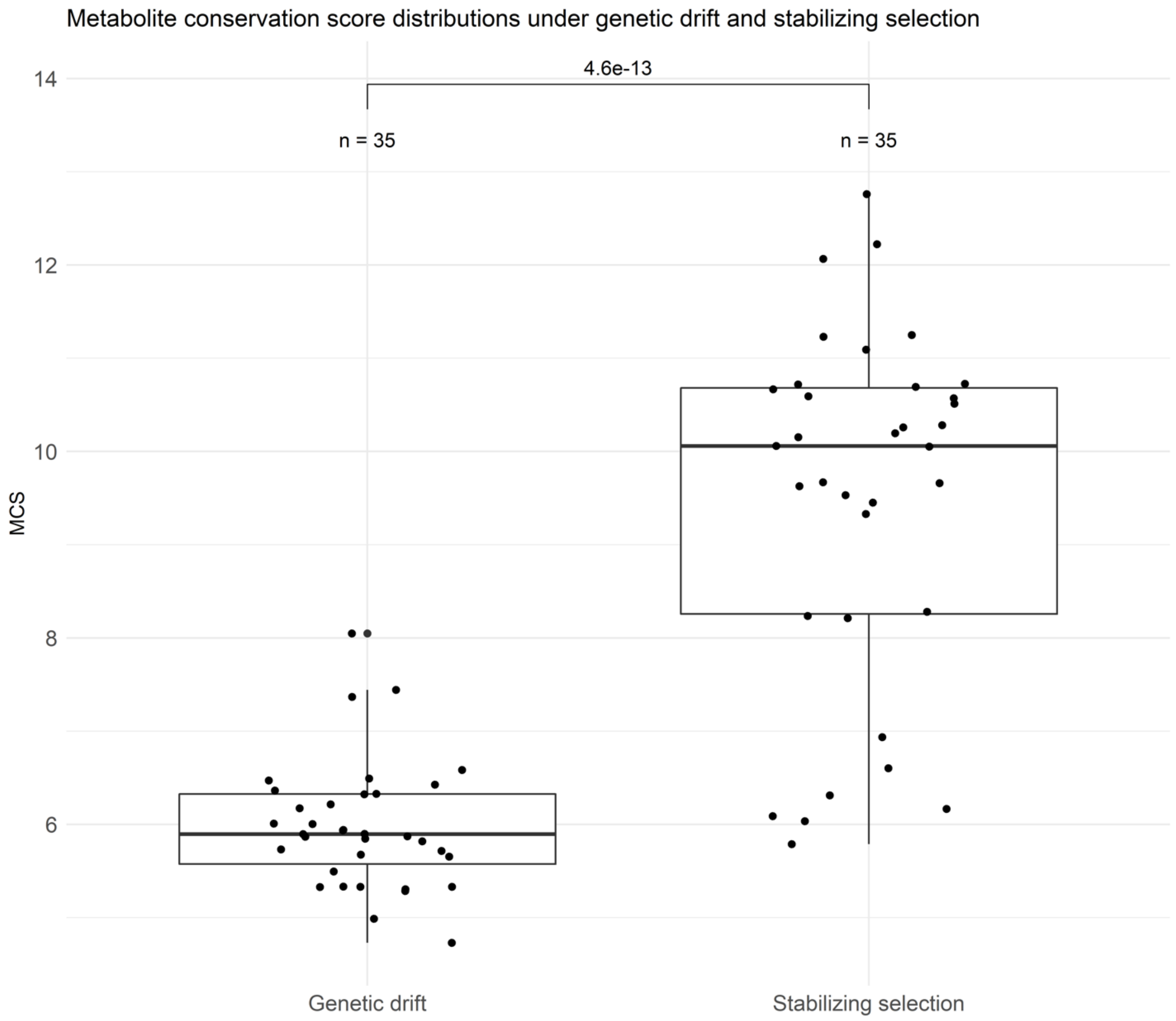


**Figure S10: Distribution of simulated metabolite conservation scores under genetic drift and stabilizing selection.** The distribution of simulated MCSs of all metabolites are shown for both selection regimes (pure genetic drift versus stabilizing selection). The median of metabolite conservation scores differ significantly between the two selection regimes (Welch two-sample t-test with unequal variance, p-value $= 4.57 \times 10^{-13}$). MCSs show larger variance under stabilizing selection as compared to pure genetic drift (F-test, F = 0.127, p $<10^{-7}$).

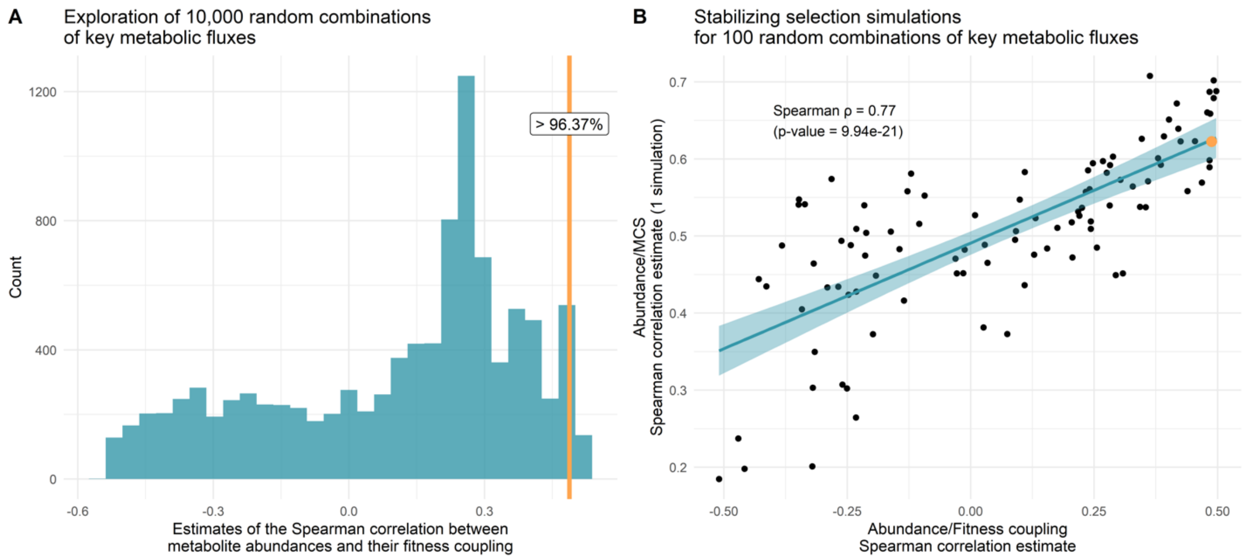


**Figure S11: The relationship between metabolite abundance and fitness coupling for randomly defined key fluxes. A.** The correlation between metabolite abundance and fitness coupling is especially strong when approximating fitness with the four key fluxes assumed to be important for erythrocyte function compared to randomly defined key fluxes. For the analysis, we calculated the fitness coupling of each metabolite to 10,000 randomly defined combinations of key fluxes (uniformly distributed from 1-uplets to 4-uplets), represented by 10,000 biochemical models. Then, the Spearman correlation is computed between metabolite abundances and their fitness coupling for each of the 10,000 model. The distribution of the 10,000 Spearman estimates is shown. The orange line represents the Spearman correlation coefficient based on the default key fluxes ($\nu_9$, $\nu_{16}$, $\nu_{21}$ and $\nu_{26}$;[8]), which is higher than 96.37% of the 10,000 random combinations. **B.** Evolutionary simulations with stabilizing selection were run for 100 metabolic models with randomly defined combinations of key metabolic fluxes (see Methods). At the end of each simulation, the Spearman correlation between metabolite abundance and *in silico* MCS was computed and plotted against the Spearman correlation between metabolite abundance and their fitness coupling calculated in the same model. The two measures show a strong positive correlation (Spearman $\rho = 0.77$, p-value $= 1.76 \times 10^{-20}$). The bold orange dot represents the estimates for the default key fluxes ($\nu_9$, $\nu_{16}$, $\nu_{21}$ and $\nu_{26}$)[8]. Thus, the correlation between metabolite abundance and *in silico* MCS is strongest in those specific biochemical models where abundant metabolites are strongly coupled to key fluxes. This results indicates that the high evolutionary conservation score of abundant metabolites is caused by their strong coupling to key fluxes in the model.

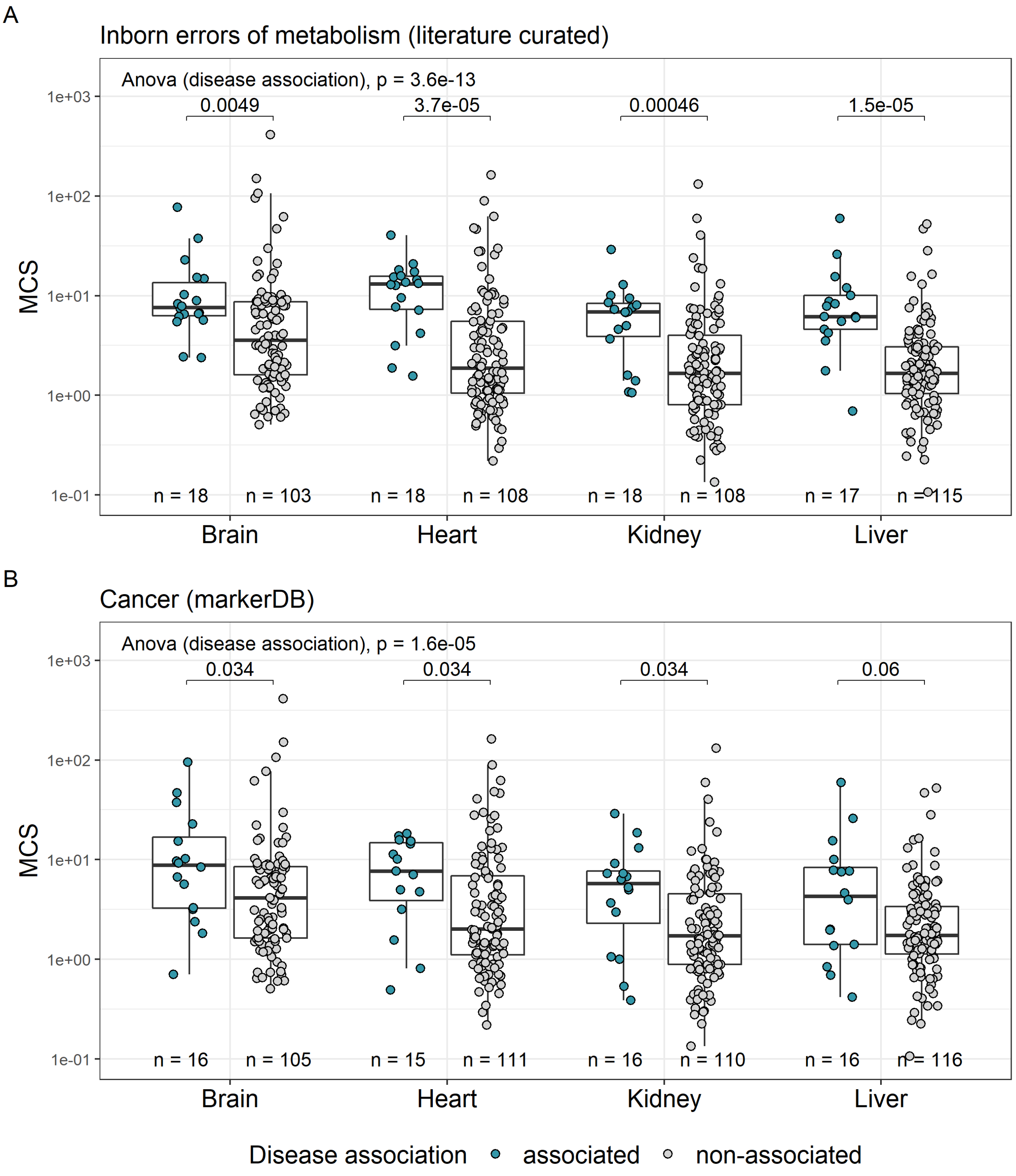
A
Inborn errors of metabolism (literature curated)
Anova (disease association), p = 3.6e-13
0.0049
3.7e-05
0.00046
1.5e-05
MCS
1e+03
1e+02
1e+01
1e+00
1e-01
n = 18
n = 103
n = 18
n = 108
n = 18
n = 108
n = 17
n = 115
Brain
Heart
Kidney
Liver
B
Cancer (markerDB)
Anova (disease association), p = 1.6e-05
0.034
0.034
0.034
0.06
MCS
n = 16
n = 105
n = 15
n = 111
n = 16
n = 110
n = 16
n = 116
Brain
Heart
Kidney
Liver
Disease association
associated
non-associated

C

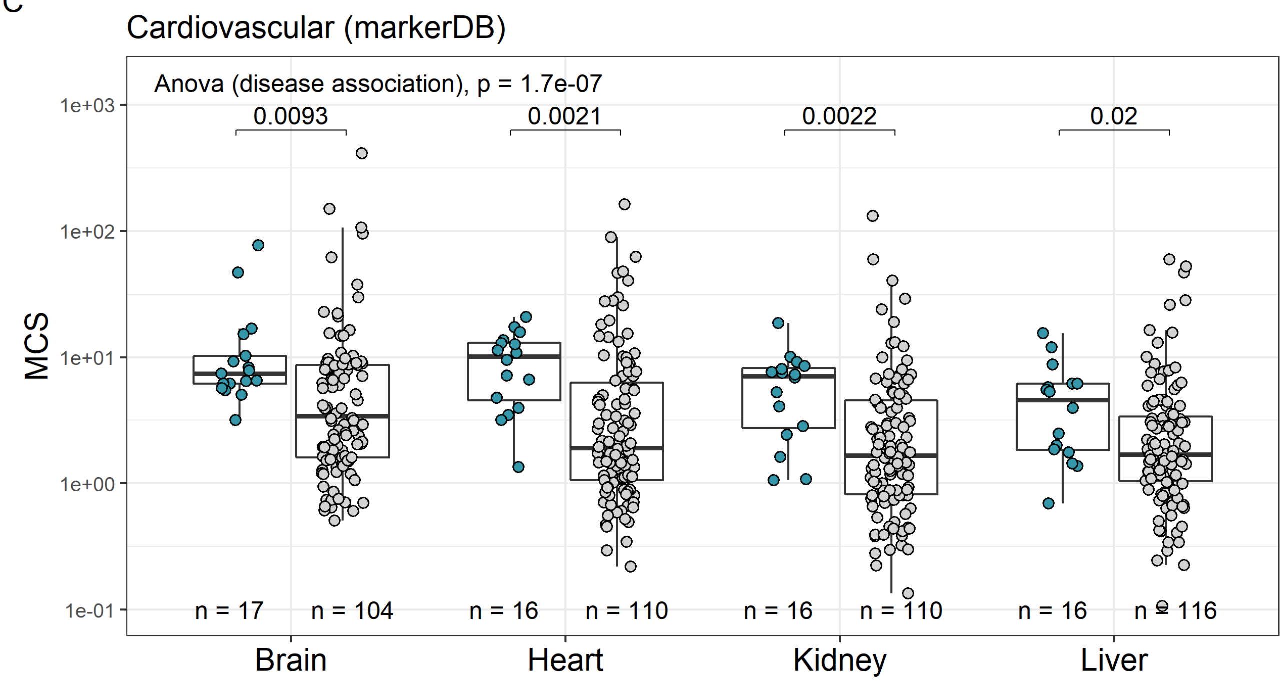


D

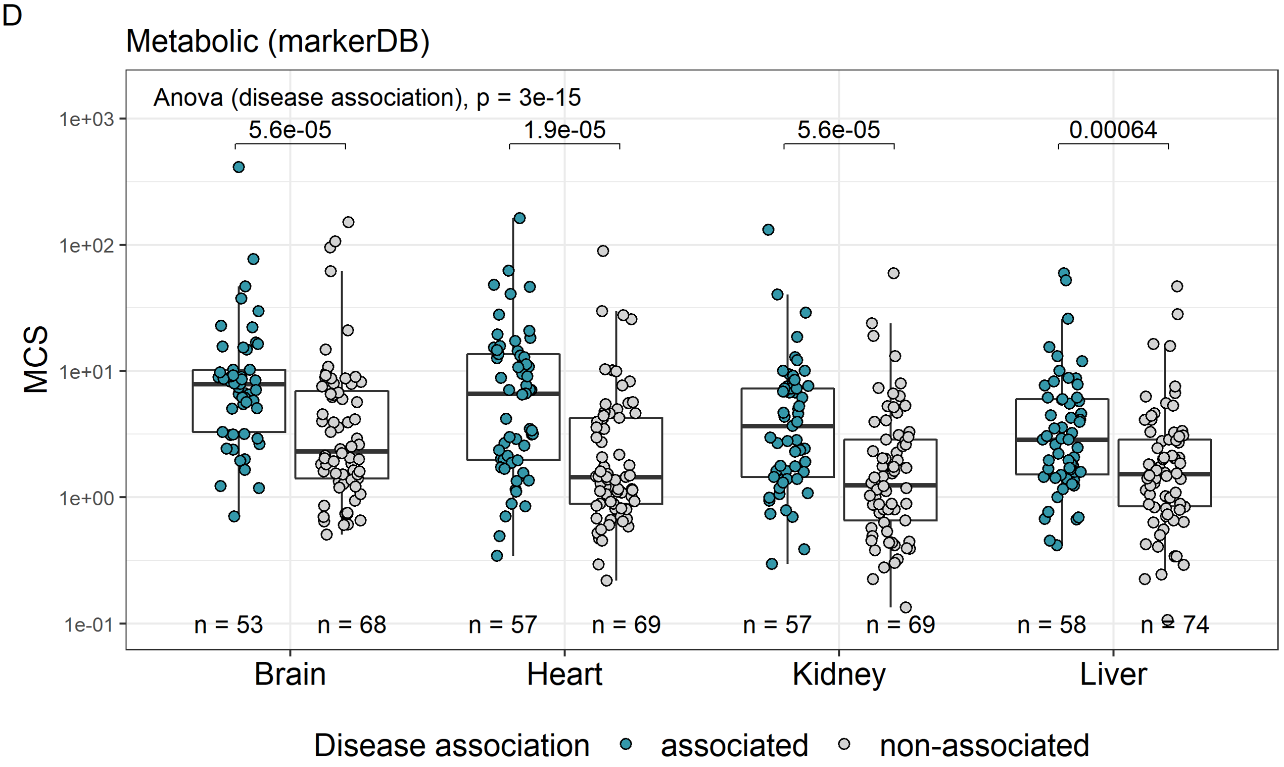


**Figure S12: Metabolites associated with certain disease conditions show high MCS in all four studied organs.** Broad disease conditions are as follows: **A**: Inborn errors of metabolism (IEM) associated metabolites according to our manual curation of the literature, **B**: cancer-associated metabolites according to MarkerDB, **C**: cardiovascular disease associated metabolites according to MarkerDB and **D**: metabolic disease associated metabolites according to MarkerDB. P-values corresponding to each individual organ are shown above the boxplots and were calculated using the two-sided Wilcoxon rank sum test.

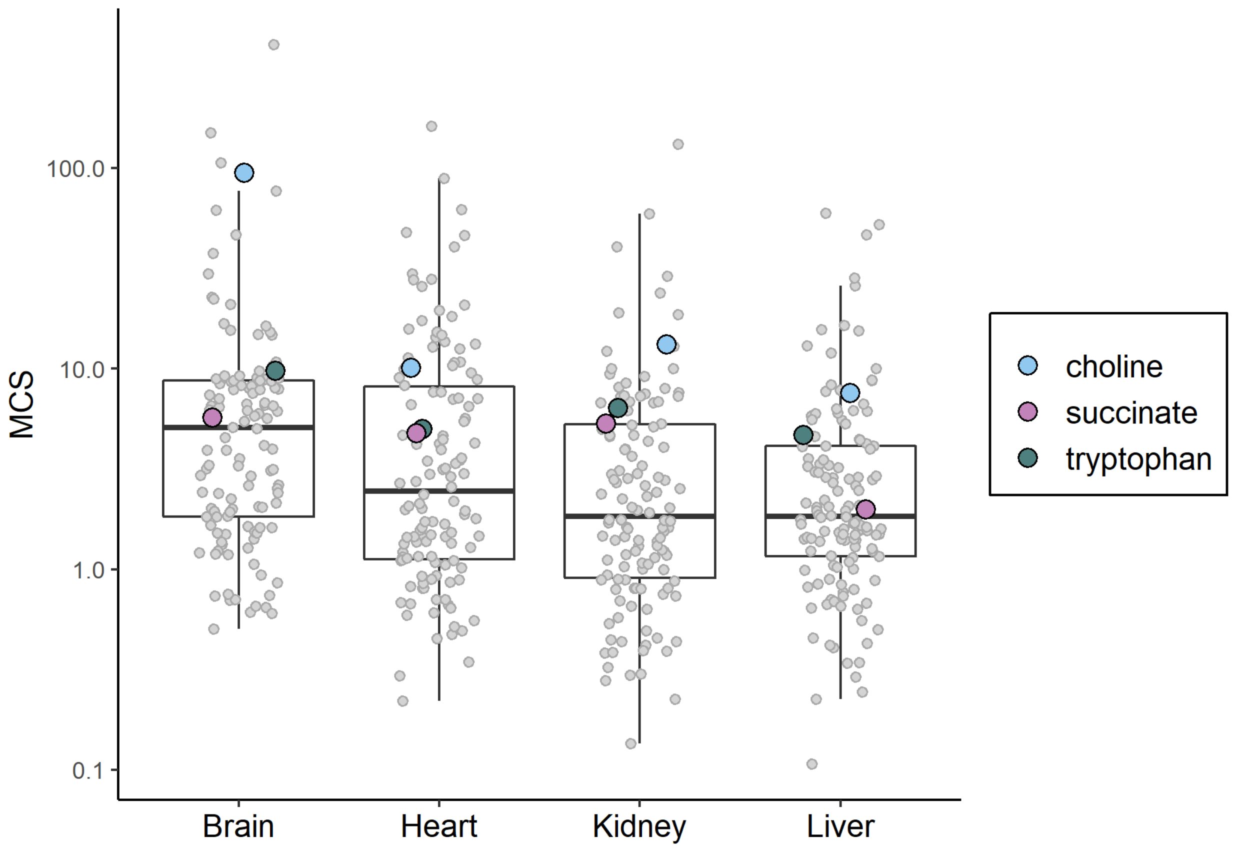


**Figure S13**: **Conservation scores of three metabolites associated with cancers that are not involved in IEMs.** Plot shows the conservation scores of all measured metabolites in the four organs (grey dots), with three specific metabolites marked with colors: choline and tryptophan are associated with several types of cancers, while succinate is a known oncometabolite and has a role in cardiovascular conditions as well. Importantly, none of the three highlighted metabolites are associated with metabolic disorders according to MarkerDB.

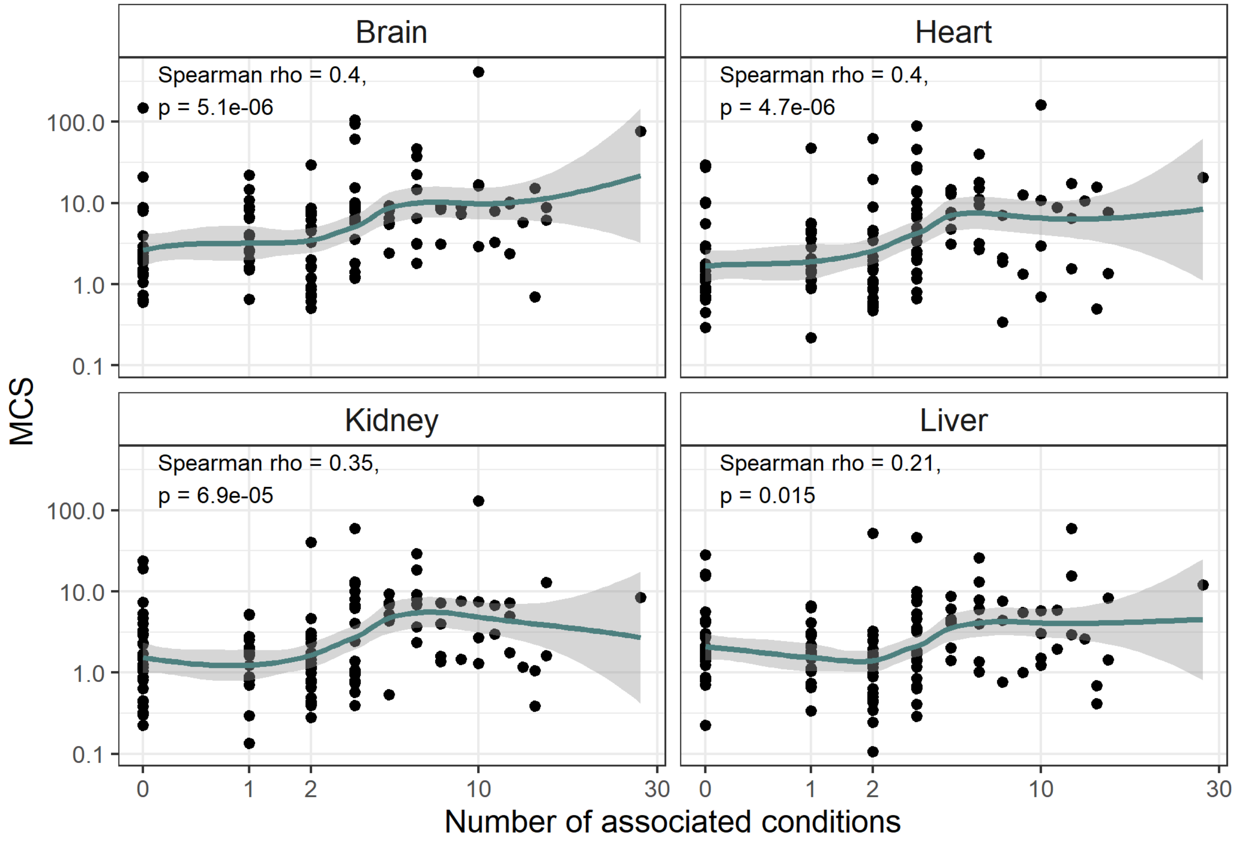


**Figure S14: Metabolites associated with many diseases display high conservation scores in mammals.** Scatterplots show the associations between conservation scores of metabolites and the number of specific disease conditions they are associated with, according to MarkerDB ($N_{Brain}$=121, $N_{Heart}$=126, $N_{Kidney}$=126, $N_{Liver}$=132). Correlations are significant in all four studied organs (Spearman's rho). Lines indicate smooth curve fitted using LOESS regression, with their 95% confidence intervals shaded in grey.

### Supplementary Tables

**Table S1: The percentage of variance in metabolite conservation score explained by individual metabolite features**. The table also shows the directions and p-values of the effects. "$R^2$ of metabolite feature" indicates the proportion of variance in MCS explained by each metabolite feature in a multiple linear regression model including both the metabolite feature and organ membership as explanatory variables to account for between-organ effects. P-value refers to significance level of the metabolite feature while accounting for organ membership. Metabolite features that have a significant effect on conservation score following false discovery rate (FDR) correction when accounting for organ membership were included in the multivariate analysis.

| Feature group | Metabolite feature name | Direction | $R^2$ of metabolite feature | FDR-adjusted p-value |
|---|---|---|---|---|
| **Essentiality** | Essentiality | positive | 0.078 | 4.57E-08 |
| **Position in metabolic network** | In biosynthetic reaction | positive | 0.028 | 0.0013 |
| | In degradation reaction | negative | 2.60E-04 | 0.75 |
| | In energy metabolism | negative | 0.0086 | 0.078 |
| **Regulation** | Is activator metabolite | positive | 0.054 | 2.39E-07 |
| | Is inhibitor metabolite | negative | 3.76E-04 | 0.68 |
| | Is co-factor | negative | 0.0081 | 0.047 |
| **Abundance** | Abundance | positive | 0.18 | 6.03E-15 |
| **Hydrophobicity** | Hydrophobicity | negative | 0.0048 | 0.13 |
| **Solubility** | Solubility | positive | 0.061 | 1.39E-07 |
| **Molecular weight** | Molecular weight | negative | 0.082 | 5.49E-10 |
| **pKa** | pKa (acidic) | negative | 0.0033 | 0.22 |
| | pKa (basic) | positive | 0.0083 | 0.060 |
| **Network degree** | Network degree | positive | 0.031 | 2.54E-04 |
| **Toxicity (LD50)** | Toxicity (LD50) | negative | 0.037 | 0.011 |
| **Metabolite class** | Aliphatic compounds | positive | 0.00015 | 0.78 |
| | Amino acids | positive | 0.024 | 0.00085 |
| | Aromatic compounds | negative | 0.014 | 0.0076 |
| | Carbohydrates | negative | 0.023 | 0.00085 |
| | Nucleosides, nucleotides | negative | 0.032 | 0.00021 |
| | Organic acids | positive | 0.023 | 0.00085 |
| | Other metabolites | positive | 0.016 | 0.0044 |
| **Pathway association (KEGG)** | TCA cycle (hsa00020) | positive | 0.0021 | 0.51 |
| | Pentose and glucoronate interconversions (hsa00040) | positive | 0.00065 | 0.69 |
| | Galactose metabolism (hsa00052) | negative | 0.0032 | 0.38 |
| | Ascorbate and aldarate metabolism (hsa00053) | positive | 0.0018 | 0.52 |

| | | | | |
|---|---|---|---|---|
| | Arginine biosynthesis (hsa00220) | positive | 0.019 | 0.017 |
| | Purine metabolism (hsa00230) | negative | 0.002 | 0.51 |
| | Pyrimidine metabolism (hsa00240) | negative | 0.0042 | 0.31 |
| | Alanine, aspartate, glutamate metabolism (hsa00250) | positive | 0.009 | 0.12 |
| | Glycine, serine, threonine metabolism (hsa00260) | positive | 0.032 | 0.0015 |
| | Cysteine, methionine metabolism (hsa00270) | positive | 0.0013 | 0.58 |
| | Valine, leucine, isoleucine biosynthesis (hsa00290) | positive | 0.026 | 0.0036 |
| | Lysine degradation (hsa00310) | positive | 0.0068 | 0.21 |
| | Arginine, proline metabolism (hsa00330) | positive | 0.0099 | 0.11 |
| | Histidine metabolism (hsa00340) | positive | 0.00079 | 0.67 |
| | Tyrosine metabolism (hsa00350) | positive | 0.0058 | 0.23 |
| | Phenylalanine metabolism (hsa00360) | positive | 0.005 | 0.27 |
| | Tryptophan metabolism (hsa00380) | negative | 0.017 | 0.02 |
| | Phenylalanine, tyrosine, tryptophan biosynthesis (hsa00400) | positive | 0.00026 | 0.76 |
| | Beta-alanine metabolism (hsa00410) | negative | 0.00032 | 0.76 |
| | Taurine, hypotaurine metabolism (hsa00430) | positive | 0.03 | 0.0015 |
| | Glutathione metabolism (hsa00480) | positive | 0.00019 | 0.77 |
| | Pyruvate metabolism (hsa00620) | positive | 0.018 | 0.018 |
| | Glyoxalate, dicarboxylate metabolism (hsa00630) | positive | 0.043 | 0.00019 |
| | Butanoate metabolism (hsa00650) | positive | 0.0012 | 0.58 |
| | Thiamine metabolism (hsa00730) | positive | 0.0063 | 0.22 |
| | Nicotinate and nicotinamide metabolism (hsa00760) | negative | 0.00025 | 0.76 |
| | Pantothenate and CoA metabolism (hsa00770) | positive | 0.004 | 0.31 |

Table S2: **Estimating the upper limit of predictability of metabolite conservation score.** To estimate the upper limit of predictability of MCS, we calculated the proportion of variance in MCS explained by MCS calculated on independent clades on the phylogenetic tree, yielding an estimate of the evolutionary reproducibility of metabolite conservation. In the table, "$R^2$ of clade-specific MCS" denotes the proportion of variance in MCS calculated in one clade explained by MCS calculated on the other clade using a multiple linear regression model including both clade-specific MCS and organ membership as explanatory variables to account for between-organ effects. P-value refers to significance level of the clade-specific MCS while accounting for organ membership.

| **Response variable** | **Explanatory variable** | **$R^2$ of clade-specific MCS** | **p-value** |
|---|---|---|---|
| MCS calculated in Clade 1 | MCS calculated in Clade 2 | 0.49 | 3.9E-82 |
| MCS calculated in Clade 2 | MCS calculated in Clade 1 | 0.48 | 3.9E-82 |

**Table S3: Statistics of the most parsimonious multivariate metabolite feature model**. The table contains the statistics of the most parsimonious multivariate model, a linear model comprised of the 7 metabolite features which best predict the extent of evolutionary conservation, as well as organ membership. The last two rows (shaded in blue) contain the independent effect of organ membership ($R^2$) and the portion of variance in MCS ($R^2$) jointly explained by the 7 metabolite features. The independent effect of organ membership was estimated by fitting a simpler multivariate model without organ membership and calculating the decrease in the adjusted $R^2$ value compared to the full multivariate model. The portion of variance explained by the metabolite features was determined by subtracting the independent effect of organ membership from the adjusted $R^2$ of the full model (Methods).

| | |
|---|---|
| *Residual standard error* | 0.46 |
| *Adjusted $R^2$ of full model* | 0.4 |
| *F- statistic* | 12.32 |
| *p-value* | 2.14e-22 |
| *Effect of organ membership ($R^2$)* | 0.091 |
| *Adjusted $R^2$ of metabolite feature effects* | 0.31 |

**Table S4: Metabolite features included in the multivariate regression models.** The table shows all the metabolite features that individually showed a significant association with conservation score (and were therefore included in the multivariate analysis). Five metabolite features and three KEGG pathways were included in the final model, following stepwise model selection. The statistical significance of these features and their independent effects in the final model are also indicated.

| Metabolite feature | Included in initial multivariate model | Included in final multivariate model | Significant in final model ($p<0.05$) | Effect of predictor ($R^2$) |
|---|---|---|---|---|
| *Abundance* | X | X | Yes | 0.12 |
| *Essentiality* | X | X | Yes | 0.033 |
| *Class* | X | X | No | 0.016 |
| *Molecular weight* | X | X | Yes | 0.040 |
| *Solubility* | X | - | - | - |
| *Network degree* | X | - | - | - |
| *Biosynthesis* | X | - | - | - |
| *Activator* | X | - | - | - |
| *Arginine biosynthesis (hsa00220)* | X | - | - | - |
| *Glycine, serine and threonine metabolism (hsa00260)* | X | X | No | 0.0062 |
| *Valine, leucine and isoleucine biosynthesis (hsa00290)* | X | X | No | 0.0023 |
| *Tryptophan metabolism (hsa00380)* | X | - | - | - |
| *Taurine and hypotaurine metabolism (hsa00430)* | X | - | - | - |
| *Pyruvate metabolism (hsa00620)* | X | - | - | - |
| *Glyoxylate and dicarboxylate metabolism (hsa00630)* | X | X | No | 0.0058 |

**Table S5: Spearman correlation between metabolite abundance and measurement error adjusted conservation scores.** The positive relationship between evolutionary conservation and metabolite abundance holds even when calculating MCS while explicitly accounting for metabolite concentration measurement variability on the species-level (based on biological replicate measurements).

| Organ | Spearman correlation rho | Spearman correlation p-value |
|---|---|---|
| Brain | 0.34 | 0.0033. |
| Heart | 0.43 | 0.00032. |
| Kidney | 0.53 | 2.023e-05. |
| Liver | 0.36 | 0.0028. |

**Table S6: Metabolites showing organ-specific conservation.** The table lists all metabolites that show the top 10% highest organ-specific deviation in conservation score in at least one organ, along with the specific organ(s) in which they are strongly conserved and their organ-specific effect size scores in brain, heart, kidney and liver. A larger deviation score indicates that a metabolite is more conserved in the given organ than in the others.

| Metabolite | Most conserved organ | Organ-specific deviation in conservation score | | | |
|---|---|---|---|---|---|
| | | Brain | Heart | Kidney | Liver |
| hypoxanthine | Brain, Heart, Kidney | 2.85 | 1.91 | 2.13 | -6.89 |
| citrate | Brain | 2.71 | -2.70 | 0.58 | -0.59 |
| trimethylamine-N-oxide | Brain | 2.60 | -0.31 | -1.31 | -0.98 |
| choline | Brain | 2.30 | -1.14 | -0.02 | -1.13 |
| DHAP/glyceraldehyde 3P | Brain | 2.29 | 0.03 | -0.82 | -1.50 |
| xanthine | Brain | 2.23 | -2.95 | 0.26 | 0.45 |
| glutamate | Brain | 2.21 | 0.35 | 0.57 | -3.13 |
| creatine | Brain, Heart | 2.13 | 2.66 | -1.71 | -3.09 |
| aconitate | Brain | 1.93 | -1.41 | -0.97 | 0.45 |
| GABA | Brain | 1.91 | -0.90 | -0.12 | -0.89 |
| arginine | Brain, Heart | 1.80 | 2.74 | -1.83 | -2.71 |
| taurine | Heart | -1.12 | 2.96 | 1.02 | -2.86 |
| inosine | Heart | 0.61 | 2.95 | -0.68 | -2.89 |
| pyroglutamic acid | Heart | -1.02 | 2.43 | 0.30 | -1.71 |
| carnitine | Heart | -0.63 | 2.18 | -0.62 | -0.93 |
| pyruvate | Heart | -0.09 | 1.96 | -1.66 | -0.20 |
| putrescine | Heart | 1.41 | 1.92 | -2.57 | -0.76 |
| phosphocholine | Heart | 0.20 | 1.60 | -1.46 | -0.34 |
| adipate | Heart | 0.77 | 1.45 | -1.56 | -0.66 |
| betaine | Kidney | -1.91 | -1.04 | 2.99 | -0.04 |
| xanthosine | Kidney | 0.18 | -2.03 | 2.23 | -0.37 |
| glucuronate | Kidney | -1.76 | -1.15 | 2.23 | 0.68 |
| hexose diphosphate | Kidney | -1.36 | -0.64 | 1.74 | 0.27 |
| adenine | Kidney | -1.21 | -0.80 | 1.50 | 0.51 |
| inositol | Kidney | 0.76 | -1.32 | 1.40 | -0.84 |
| cystathionine | Kidney | -1.01 | -0.79 | 1.25 | 0.56 |
| glycine | Kidney | -1.03 | -0.44 | 1.18 | 0.28 |
| aspartate | Kidney | -0.96 | 0.54 | 1.09 | -0.66 |
| citrulline | Kidney | -1.65 | -0.28 | 1.07 | 0.86 |
| fructose/glucose/galactose | Liver | -2.64 | -2.59 | 0.26 | 4.98 |
| uracil | Liver | -1.42 | -0.86 | -0.47 | 2.75 |
| 3-phosphoglycerate | Liver | -1.33 | 0.16 | -1.16 | 2.33 |
| hexose monophosphate | Liver | -1.46 | -0.28 | -0.40 | 2.13 |
| phosphoenolpyruvate | Liver | -0.14 | -0.05 | -1.64 | 1.82 |

| | | | | | |
|---|---|---|---|---|---|
| kynurenine | Liver | -0.96 | -0.53 | -0.24 | 1.73 |
| NADP | Liver | -1.33 | -0.92 | 0.59 | 1.65 |
| 4-pyridoxate | Liver | -1.76 | -0.30 | 0.47 | 1.59 |
| lactose | Liver | 0.43 | -0.21 | -1.70 | 1.48 |
| dUMP | Liver | -1.37 | -0.01 | -0.05 | 1.44 |
| GDP | Liver | -0.11 | -2.02 | 0.71 | 1.41 |

**Table S7: Independent effects of metabolite abundance and reaction essentiality on metabolite conservation score in the evolutionary simulations.**

Here we tested whether two metabolite properties, metabolite abundance and involvement in essential reaction, are independently associated with metabolite conservation scores in the presence of stabilizing selection in the erythrocyte metabolic model. For 32 metabolites, metabolite abundance and involvement in essential reaction were calculated from biochemical modelling of the wild-type erythrocyte network, while metabolite conservation scores were calculated from evolutionary simulations (see Methods). We used multivariate linear modelling to predict metabolite conservation score based on metabolite abundance and involvement in essential reaction. Note that metabolite conservation scores and metabolite abundances were log-transformed, while involvement in essential reaction was a binary variable. A linear model was fitted using the R function lm() and explained well variation in conservation scores across metabolites (adjusted R-squared: 0.65, p-value = 7.70e-08). Both involvement in essential reaction and metabolite abundance showed a highly significant effect in the multivariate model, indicating that these properties independently determine metabolite conservation score.

| **Explanatory variable** | **Estimate** | **Standard error** | **t-value** | **P-value** | |
|---|---|---|---|---|---|
| (Intercept) | 10.12 | 0.49 | 21.57 | < 2e-16 | *** |
| In essential reaction | 1.85 | 0.42 | 4.35 | 1.55e-04 | *** |
| Metabolite abundance | 0.81 | 0.15 | 5.23 | 1.35e-05 | *** |

**Table S8: Metabolites associated with inborn errors of metabolism.** Only the metabolites present in our evolutionary dataset are shown.

| Disease name | OMIM ID | Orphanet ORPHAcode | Causative gene name | Gene NCBI ID | Associated metabolite name | Associated metabolite HMDB ID |
|---|---|---|---|---|---|---|
| *Propionic Acidemia* | 606054 | ORPHA:35 | PCCA; PCCB | 5095; 5096 | propionate | HMDB0000237 |
| *Methylmalonic Acidemia (methylmalonyl-CoA mutase)* | 251000 | ORPHA:27; ORPHA:79312; ORPHA:289916 | MMUT | 4594 | propionate | HMDB0000237 |
| *Methylmalonic Acidemia (Cobalamin disorders)* | 277410; 251100; 251110; 251120 | ORPHA:26; ORPHA:28; ORPHA:622; ORPHA:79283; ORPHA:79310; ORPHA:79311; ORPHA:308380; ORPHA:308442; ORPHA:308425 | MMAA; MMAB ;MMADHC; MCEE | 166785; 326625; 27249; 84693 | propionate | HMDB0000237 |
| *Isovaleric Acidemia* | 243500 | ORPHA:33 | IVD | 3712 | | |
| *3-Methylcrotonyl-CoA Carboxylase Deficiency* | 210200; 210210 | ORPHA:6 | MCCC1; MCCC2 | 56922; 64087 | | |
| *3-Hydroxy-3-Methyglutaric Aciduria* | 246450 | ORPHA:20 | HMGCL | 3155 | | |
| *Holocarboxylase Synthase Deficiency* | 253270 | ORPHA:79242 | HLCS | 3141 | lactate | HMDB0000190 |
| *beta-Ketothiolase Deficiency* | 203750 | ORPHA:134 | ACAT1 | 38 | | |
| *Glutaric acidemia* | 231670 | ORPHA:25 | GCDH | 2639 | | |
| *Carnitine Uptake Defect/Carnitine Transport Defect* | 212140 | ORPHA:158 | SLC22A5 | 6584 | carnitine | HMDB0000062 |
| *Medium-chain Acyl-CoA Dehydrogenase Deficiency* | 201450 | ORPHA:42 | ACADM | 34 | | |
| *Very Long-chain Acyl-CoA Dehydrogenase Deficiency* | 201475 | ORPHA:26793 | ACADVL | 37 | | |
| *Long-chain L-3 Hydroxyacyl-CoA Dehydrogenase Deficiency* | 609016 | ORPHA:5 | HADHA | 3030 | suberate | HMDB0000893 |
| | | | | | adipate | HMDB0000448 |
| *Trifunctional Protein Deficiency* | 609015 | ORPHA:746 | HADHA; HADHB | 3030; 3032 | suberate | HMDB0000893 |
| | | | | | adipate | HMDB0000448 |
| *Argininosuccinic Aciduria* | 207900 | ORPHA:23 | ASL | 435 | alanine | HMDB0000161 |
| | | | | | citrulline | HMDB0000904 |
| | | | | | glutamine | HMDB0000641 |
| | | | | | glycine | HMDB0000123 |

| | | | | | | |
|---|---|---|---|---|---|---|
| *Citrullinemia, Type I* | 215700 | ORPHA:247525 | ASS1 | 445 | citrulline | HMDB0000904 |
| | | | | | orotate | HMDB0000226 |
| *Maple Syrup Urine Disease* | 248600 | ORPHA:511; ORPHA:268145; ORPHA:268162; ORPHA:268173; ORPHA:268184 | BCKDHA; BCKDHB; DBT | 593; 594; 1629 | fumarate/maleate /alpha-ketoisovalerate | HMDB0000019 |
| | | | | | isoleucine | HMDB0000172 |
| | | | | | leucine | HMDB0000687 |
| | | | | | valine | HMDB0000883 |
| *Homocystinuria* | 236200; 236250; 236270; 250940 | ORPHA:394; ORPHA:395; ORPHA:622; ORPHA:2169; ORPHA:2170 | CBS; MTHFR; MTR; MTRR | 875; 4524; 4548; 4552 | methionine | HMDB0000696 |
| *Classic Phenylketonuria* | 261600 | ORPHA:716; ORPHA:2209 | PAH | 5053 | phenylalanine | HMDB0000159 |
| *Tyrosinemia, Type I* | 276700 | ORPHA:882 | FAH | 2184 | tyrosine | HMDB0000158 |
| *Biotinidase Deficiency* | 253260 | ORPHA:79241 | BTD | 686 | | |
| *Classic Galactosemia* | 230400 | ORPHA:352; ORPHA:79239 | GALT | 2592 | fructose/glucose/ galactose | HMDB0000143 |
| *Glycogen Storage Disease Type II (Pompe disease)* | 232300 | ORPHA:365 | GAA | 2548 | | |
| *Mucopolysaccharidosis Type 1* | 607014; 607015; 607016 | ORPHA:579; ORPHA:93473; ORPHA:93474; ORPHA:93476 | IDUA | 3425 | | |

**Table S9: Metabolites associated with inborn errors of metabolism show elevated conservation scores independent of amino acid and metabolic pathway overrepresentations and the effect of metabolite abundance.**

We tested whether IEM-associated metabolites have higher metabolite conservation scores than the rest of metabolites under four scenarios: (i) for the entire dataset, (ii) when removing all amino acids from the dataset, (iii) when removing metabolites involved in one of the pathways containing three or more IEM-associated metabolites, namely *Arginine biosynthesis* (hsa00220) and *Valine, leucine and isoleucine biosynthesis* (hsa00290), and (iv) when controlling for the effect of metabolite abundance. In the first three cases, we used ANOVA models accounting for both organ membership and IEM-association, while the effect of metabolite abundance was controlled for by building a linear model with both abundance and IEM-association as explanatory variables. IEM-associated metabolites were significantly more conserved in all cases, and both abundance and IEM-association had significant independent effects on metabolite conservation score.

| **Test** | **Explanatory variable** | **Effect of IEM-association** | | |
|---|---|---|---|---|
| | | **F-value** | **p-value** | |
| All IEM associations | IEM-association | 55.82 | 3.6e-13 | *** |
| No amino acids | IEM-association | 25.29 | 8e-07 | *** |
| No *hsa00220* metabolites | IEM-association | 54.4 | 7.07e-13 | *** |
| No *hsa00290* metabolites | IEM-association | 44.47 | 7.03e-11 | *** |
| Corrected for abundance | IEM association | 37.6 | 2.7e-09 | *** |
| | Abundance | 20.79 | 7.4e-06 | *** |

**Table S10: Testing the associations between MarkerDB disorder categories and metabolite conservation in a multivariate model**. We used a multivariate linear regression model to identify broad disorder categories whose biomarkers show an elevated conservation score compared to the rest of metabolites. Note that only disorder categories containing at least ten metabolites from our dataset and showing a significant association with conservation score (single-variable two-sided Wilcoxon rank sum test $p<0.05$) in at least three organs were included in the multivariate linear regression model.

| Disease category | # of metabolites | # of significant tissues | Included in the multivariate model | Significant relationship between MCS and disorder category in the multivariate model ($p<0.05$) |
|---|---|---|---|---|
| *Cancer* | 16 | 3 | X | Yes |
| *Cardiovascular system disorder* | 17 | 3 | X | Yes |
| *Digestive system disorder* | 33 | 2 | - | - |
| *Endocrine system disorder* | 16 | 0 | - | - |
| *Germline disorder* | 71 | 3 | X | No |
| *Hematological or lymphatic system disorder* | 16 | 1 | - | - |
| *Immune system disorder* | 38 | 1 | - | - |
| *Mental or behavioral disorder* | 56 | 2 | - | - |
| *Metabolic disorder* | 60 | 4 | X | Yes |
| *Nervous system disorder* | 74 | 0 | - | - |
| *Urinary system disorder* | 32 | 1 | - | - |

**Table S11: Metabolites associated with many diseases are more conserved, even when accounting for metabolite abundance and essentiality.** Statistics of two linear models that assess the relationship between the conservation score of metabolites and the number of human diseases they are associated with while accounting for (A) metabolite abundance and (B) metabolite essentiality. Metabolites associated with a larger number of human diseases are significantly more conserved even after controlling for the effects of metabolite abundance and essentiality. Both linear models used also included organ membership as an explanatory variable, so that the effects of disease participation and metabolite features could be assessed for all organs in a single model.

| **Test** | **Variable** | **F-value** | **P-value** | |
|---|---|---|---|---|
| *A: Controlling for abundance* | Number of associated disorders | 21.93 | 4.5e-06 | *** |
| | Metabolite abundance | 49.52 | 1.8e-11 | *** |
| *B: Controlling for essentiality* | Number of associated disorders | 23.46 | 2.02e-06 | *** |
| | Involvement in essential reactions | 18.42 | 2.4e-05 | *** |

**Table S12: List of simulation parameters for each *in silico* evolution experiment.**

| Parameter name | Symbol | Genetic drift simulations | Stabilizing selection simulations |
|---|---|---|---|
| Selection threshold | $\omega$ | $+\infty$ | $1 \times 10^{-4}$ |
| Mutation size | $\sigma_{mut}$ | $1 \times 10^{-2}$ | |
| Number of iterations | $T$ | $10{,}000$ | |
| Repetitions | - | $10$ | |

**Appendix S1: Evaluation of two additional kinetic models**

To further test whether stabilizing selection on key metabolic fluxes explains patterns of metabolite conservation (see main text), we performed the same analyses as for the erythrocyte metabolism model[1] for two other publicly available kinetic models: another model of human erythrocyte metabolism[2] (56 variables metabolites, 53 reactions, 234 kinetic parameters) and a model of human hepatic glucose metabolism[3] (24 variable metabolites, 36 reactions, 200 kinetic parameters). The mutation size was $\sigma_{mut} = 1 \times 10^{-2}$ for all the simulations and fitness coupling analyses; the selection threshold was $\omega = 1 \times 10^{-5}$ for the erythrocyte model and $\omega = 1 \times 10^{-2}$ for the hepatocyte model. We ran the evolutionary simulations for $T = 10{,}000$ iterations (10 repetitions per experiment). Five low-varying metabolites (glutathione, NAD, NADPH, phosphate, and pyruvate) have been removed in the erythrocyte model, based on the filtering approach described in Methods. One low-varying metabolite (mitochondrial GTP) has been removed in the hepatocyte model. For the erythrocyte model, the key metabolic fluxes were ATP utilization (*vatpase*), glutathione (GSH) oxidation (*vox*) and the formation of 2,3-bisphosphoglycerate (*vbpgsp7*). For the hepatocyte model, the key metabolic fluxes were the reactions associated with blood glucose regulation: glycogenesis (fluxes *GS* and *GP*), the production of oxaloacetate from pyruvate, a precursor of gluconeogenesis (*PC*) and the production of Acetyl-CoA, a precursor of triglycerides (*PDH*).

For each kinetic model, we also computed the fitness coupling of metabolites for 10,000 random combinations of key fluxes, and we computed stabilizing selection simulations for 100 random combinations of key fluxes (from 1-uplets up to 3-uplets for the erythrocyte model; from 1-uplets to 4-uplets for the hepatocyte model). We also computed a flux inhibition analysis for each model to determine essential reactions and associated metabolites (see Methods). Fluxes were constrained to 1% of their wild-type levels for the erythrocyte model (numerical instability for flux values close to zero did not allow us to use lower values). Fluxes were constrained to 0.001% for the hepatocyte model.

We also found that metabolite abundances positively correlate with their conservation scores under stabilizing selection, but not under genetic drift (Figs App. 1A-B, App. 2A-B), and that abundant metabolites are significantly more coupled to fitness (Figs App. 1C, App. 2C). Essential metabolites have significantly higher conservation scores compared to non-essential metabolites under stabilizing selection, but not under genetic drift (Figs. App. 1D-E, App. 2D-E).

Furthermore, for both models, a linear regression model shows that both metabolite abundance and metabolite essentiality are independent predictors of *in silico* metabolite conservation scores under stabilizing selection (Tables App. 1 and App. 2). Similarily to the first erythrocyte model[1], the key metabolic fluxes generally assumed to be important for the fitness are significantly more coupled to abundant metabolites than 99% of 10,000 random combinations of key fluxes for both additional models (Figs App. 1F, App. 2F). Moreover, when evolution was simulated under stabilizing selection for 100 models with randomly defined key fluxes, we again found that metabolite conservation score correlates more strongly with abundance in those models in which abundant metabolites are strongly coupled to fitness (Figs App. 1G, App. 2G).

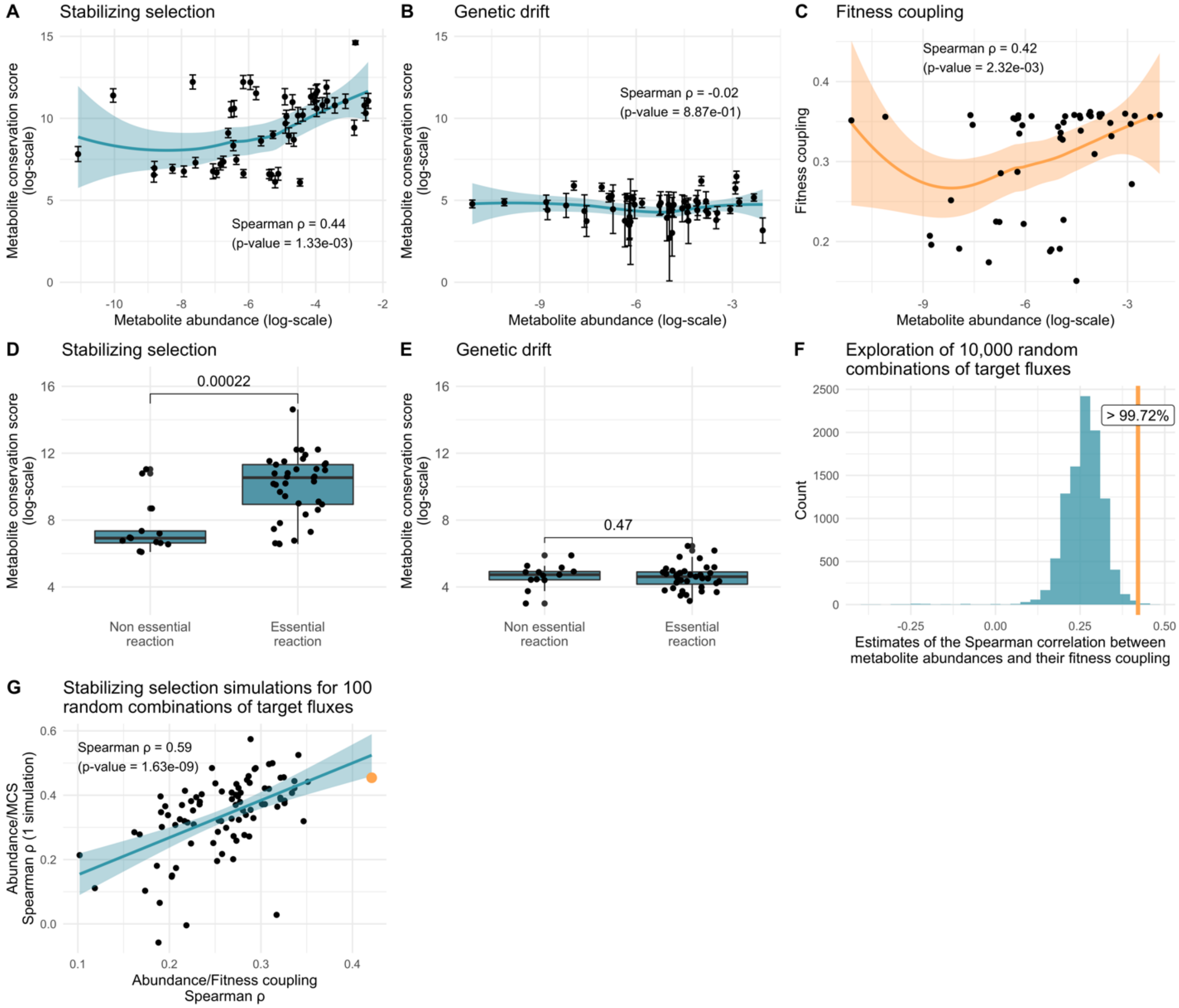


**Figure App. 1: Functional constraints in the human erythrocyte metabolism model**[2]**. A-B:** Metabolites with a higher abundance in the wild-type model show higher conservation scores in the presence of stabilizing selection (panel A, Spearman $\rho = 0.44$, $p = 1.33 \times 10^{-3}$, $N = 51$), but not in the absence of selection (panel B, Spearman $\rho = -0.02$, $p = 0.887$, $N = 51$). Each dot and error bar represents the mean and standard deviation of metabolite conservation scores of a particular metabolite based on 10 simulations. Blue lines represent LOESS regressions, with their 95% confidence interval in shaded blue. **C:** Wild-type abundances of metabolites correlate with their extent of fitness coupling (Spearman $\rho = 0.49$, $p = 3.14 \times 10^{-3}$). The orange line represents a LOESS regression, with its 95% confidence interval in shaded orange. **D-E:** Metabolites involved in essential reactions show higher conservation scores than those involved in non-essential reactions in the presence of stabilizing selection (panel D, two-sided Wilcoxon test $p =$

$2.22 \times 10^{-4}$), but not in the absence of selection (panel E, two-sided Wilcoxon test $p = 0.393$). Boxplots show the median, first and third quartiles, with the whiskers showing the values within a 1.5 inter-quartile range distance from the 1st and 3rd quartiles. Each dot represents the mean conservation score of a particular metabolite based on 10 simulations. Involved metabolites are reactants and products of the reaction (see Methods) **F:** The fitness coupling of metabolite abundances has been computed for 10,000 random combinations of key fluxes (uniformly distributed from 1-uplets to 3-uplets). Then, the Spearman correlation is computed between metabolite abundance and fitness coupling across metabolites. The distribution of the 10,000 Spearman estimates is shown. The orange line represents the Spearman estimate of the default key fluxes (vatpase, vox, vbpgsp7), which is higher than 99.72% of the 10,000 random combinations. **G:** Stabilizing selection simulations have been run for 100 random combinations of key fluxes. At the end of each simulation, an *in silico* conservation score was calculated for each metabolite. Then, for each simulation, we plotted the Spearman correlation estimate between metabolite abundance and conservation score across metabolites against the Spearman estimate of the correlation between metabolite abundance and fitness coupling across metabolites. The two Spearman estimates are significantly positively correlated (Spearman $\rho = 0.59$, $p = 1.63 \times 10^{-9}$). The bold orange dot represents the Spearman estimates for the default key fluxes (vatpase, vox, vbpgsp7).

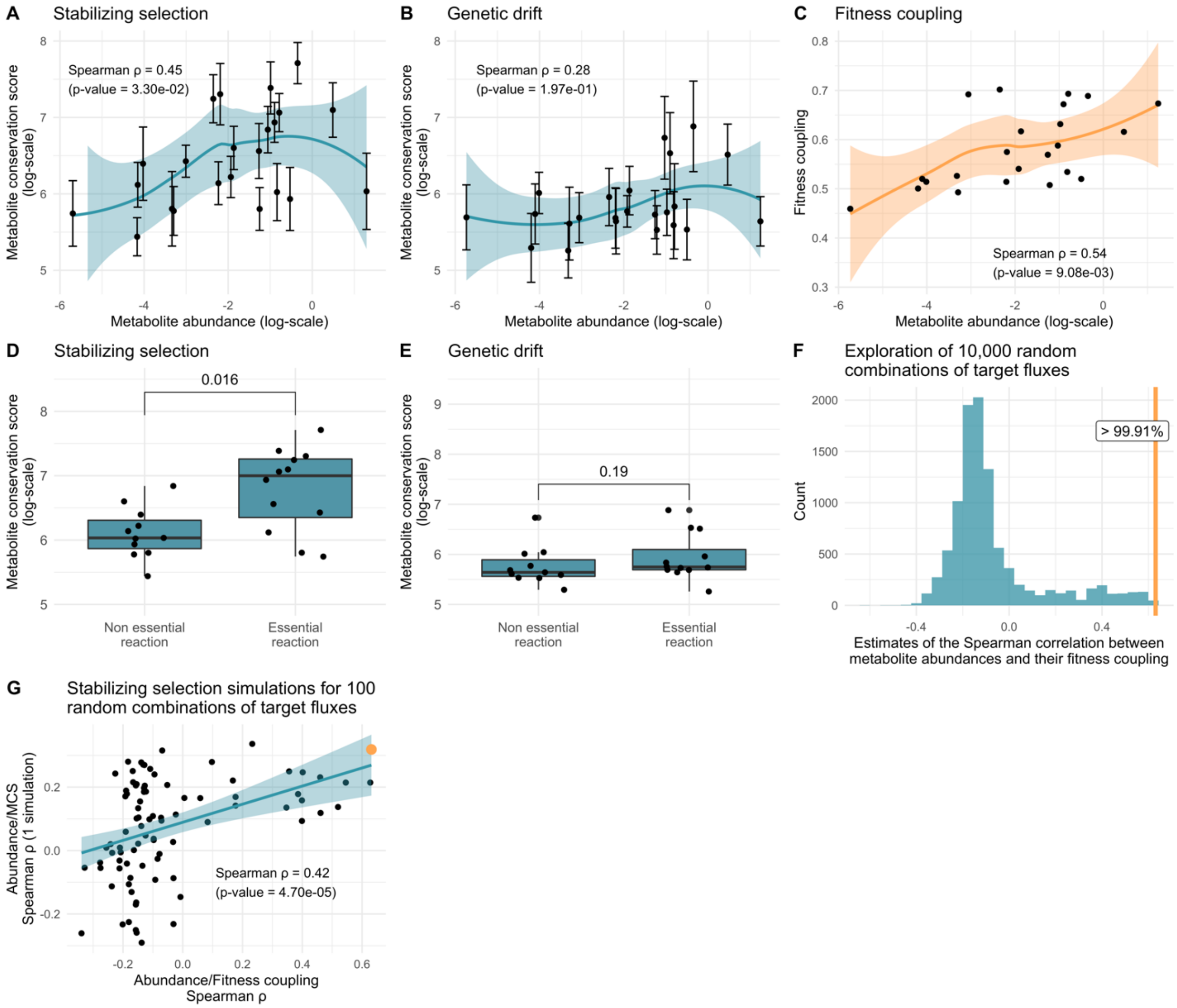


**Figure App. 2: Functional constraints in the human hepatic glucose metabolism model[3].** **A-B:** Metabolites with a higher abundance in the wild-type model show higher conservation scores in the presence of stabilizing selection (panel A, Spearman $\rho = 0.45$, p $= 3.30 \times 10^{-2}$, $N = 23$), but not in the absence of selection (panel B, Spearman $\rho = 0.28$, p $= 0.197$, $N = 23$). Each dot and error bar represents the mean and standard deviation of evolutionary rate of a particular metabolite based on 10 simulations. Blue lines represent LOESS regressions, with their 95% confidence interval in shaded blue. **C:** Wild-type abundances of metabolites correlate with their extent of fitness coupling (Spearman $\rho = 0.52$, p $= 1.08 \times 10^{-2}$). The orange line represents a LOESS regression, with its 95% confidence interval in shaded orange. **D-E:** Metabolites involved in essential reactions show higher conservation scores than those involved in non-essential reactions in the presence of stabilizing selection (panel D, two-sided Wilcoxon test p $= 0.016$), but not in the absence of selection (panel E, two-sided Wilcoxon test p $= 0.190$). Boxplots show the median, first and third quartiles, with the whiskers showing the values within a 1.5 inter-quartile range distance from the $1^{st}$ and $3^{rd}$ quartiles. Each dot represents the mean conservation score of a particular metabolite based on 10 simulations. Involved metabolites are reactants and products of the reaction (see Methods)**. F:** The fitness coupling of metabolite abundances has been computed for 10,000 random combinations of key fluxes (uniformly distributed from 1-uplets to 4-uplets). Then, the Spearman correlation is computed between metabolite abundances and their fitness coupling. The distribution of the 10,000 Spearman estimates is shown. The orange line represents the Spearman estimate of the default key fluxes (*GS, GP, PC, PDH*), which is higher than 99.91% of the 10,000 random combinations. **G.** Stabilizing selection simulations have been run for 100 random combinations of key fluxes. At the end of each simulation, an *in silico* conservation score was calculated for each metabolite. Then, for each simulation, we plotted the Spearman correlation estimate between metabolite abundance and conservation score across metabolites against the Spearman estimate of the correlation between metabolite abundance and fitness coupling across metabolites. The two Spearman estimates are significantly positively correlated (Spearman $\rho = 0.42$, p $= 4.53 \times 10^{-5}$). The bold orange dot represents the Spearman estimates for the default key fluxes (*GS, GP, PC, PDH*).

**Table App. 1: For the erythrocyte model[2], a linear regression model shows that metabolite abundance and metabolite essentiality are predictors of *in silico* metabolite conservation score under stabilizing selection.** The function lm (R 4.0.0) is used with no interaction terms on log-transformed metabolite conservation scores and metabolite abundances (metabolite essentiality being a categorical variable). Adjusted R-squared: 0.3701, p-value < 7.203e-06 (p-value stars: "*": p < 0.05, "**": p < 0.01, "***": p < 0.001).

| Explanatory variable | Estimate | Standard error | t value | P-value | |
|---|---|---|---|---|---|
| (Intercept) | 9.9651 | 0.9171 | 10.866 | 2.05e-14 | *** |
| Metabolite essentiality | 2.1493 | 0.5579 | 3.852 | 0.000354 | *** |
| Metabolite abundance | 0.3939 | 0.1272 | 3.098 | 0.003289 | ** |

**Table App. 2: For the hepatocyte model[3], a linear regression model shows that metabolite abundance and metabolite essentiality are predictors of *in silico* metabolite conservation score under stabilizing selection.** The function lm (R 4.0.0) is used with no interaction terms on log-transformed metabolite conservation scores and metabolite abundances (metabolite essentiality being a categorical variable). Adjusted R-squared: 0.4877, p-value = 4.81e-4 (p-value stars: "*": $p < 0.05$, "**": $p < 0.01$, "***": $p < 0.001$).

| Explanatory variable | Estimate | Standard error | t value | P-value | |
|---|---|---|---|---|---|
| (Intercept) | 6.44448 | 0.17188 | 37.495 | < 2e-16 | *** |
| Metabolite essentiality | 0.71701 | 0.18879 | 3.798 | 0.00113 | ** |
| Metabolite abundance | 0.18502 | 0.05809 | 3.185 | 0.00465 | ** |
